# Integrative Modelling of Innate Immune Response Dynamics during Virus Infection

**DOI:** 10.1101/2025.06.17.660089

**Authors:** Ramya Boddepalli, Harsh Chhajer, Rahul Roy

## Abstract

Positive-sense RNA viruses that constitute a large class of human pathogens employ various strategies to suppress and evade host immune defenses. Understanding the dynamic interaction between the viral life cycle and immune signaling is crucial to designing effective antiviral strategies. Although significant progress has been made, quantitative models that can accurately capture the intricate interactions and the intertwined dynamics during viral infection of cells remain missing. In this study, we develop a comprehensive mathematical model that integrates the intracellular viral life cycle with key cellular innate immune pathways, including RIG-I-mediated detection and JAK-STAT signaling. The model provides mechanistic insights into long-standing observations, capturing both virus-specific dynamics and innate immune response, and the key components driving their coupled dynamics. For example, a comparison of viruses shows how the Japanese Encephalitis virus undergoes a dramatic reduction in viral load in cells, due to its rapid replication that robustly activates the RIG-I pathway, in contrast to the poor immune control of HCV. More importantly, our model demonstrates how virus-host interactions exhibit a sharp bifurcation behavior, where minor differences in immune strength or viral suppression capacity can determine whether infections resolve or persist. We propose that ISG mRNA translation and viral replication predominantly dictate these bimodal infection outcomes. Additionally, the model not only recapitulates but also highlights molecular players involved in IFN desensitization. We demonstrate how our model’s ability to capture IFN dynamics allows us to predict optimal timing and dosing strategies for interferon-based prophylactic therapies. Together, our approach reveals fundamental features that govern the delicate balance between the establishment of infection and immune control in RNA virus infections.

**Author Summary:** Viruses responsible for diseases like hepatitis and dengue rapidly proliferate by invading host cells, which in turn activate immune responses to counteract the infection. Viruses have also developed a variety of mechanisms to interfere with this immune response, thereby complicating the process of effectively treating infections. Here, we simulate the comprehensive sequence of molecular-level actions and the corresponding counteractions that occur between the virus and the host cell components as the infection proceeds. We find that the battle between the virus and the immune system behaves like a seesaw, and small changes can dramatically tip the balance. Sometimes, a tiny boost to our immune response or a small weakness in the virus can mean the difference between cells succumbing to infection or recovering completely. Similarly, it is now possible to recognize why specific viruses are more effectively controlled by the immune system compared to others. We can now explain the decline in the immune response to interferon therapy with prolonged use and suggest the optimal timing and dosing for such treatments.

For example, we found that interferon treatment administered before infection, even in small doses, can block the virus more effectively than treatment after infection begins. In addition, we propose better ways to inhibit the virus by effectively combining complementary approaches, such as blocking its replication or its ability to hide from the immune system, which can boost the effectiveness of interferon treatment. This work provides a roadmap for developing quantitative and more effective antiviral approaches by understanding the precise biological mechanisms that determine how hosts can overcome virus infection.

## Introduction

Positive-sense RNA viruses infect hundreds of millions of people annually, leading to widespread morbidity and mortality, and constitute a significant global health burden. Since 2019, COVID-19 has been responsible for *∼*700 million cases and 7 million deaths [1]. Similarly, dengue virus accounted for *∼*6.5 million cases and *∼*7,000 deaths in 2023 [2], while Hepatitis C virus continues to cause over 1 million new infections and *∼*242,000 deaths each year [3]. This immense global impact stems from a lack of potent antiviral therapies and poor efficacy of vaccines. Positive-sense RNA ((+)RNA) viruses leverage their ability to hijack host cells, where they manipulate cellular machinery to propagate and evade immune defenses, driving a complex interplay between the virus and the host cell that underpins their pathogenicity. Although experimental approaches have identified many key host and viral molecules and their roles in defining the extent of infection in cells, our understanding of how viruses can overcome host cell immune responses remains limited and fragmented. Developing a deep and quantitative understanding of this cellular-level host-virus dynamics could be key to crafting effective antiviral strategies in the future.

The (+)RNA viral life cycle within the immune cell encompasses several coordinated, interconnected, and counteracting steps. Virus growth proceeds through cell receptor-mediated entry, host cell-assisted viral RNA translation, compartmentalized replication, coordinated assembly, and processing and release of new virions [4, 5, 6]. The intricate interplay between viral and host molecular processes is central to the outcome of infection. For instance, viral RNA translation by host cell ribosomes generates structural proteins for viral assembly and non-structural proteins that drive replication and manipulate host cellular processes to create a conducive environment for viral growth. Viral replication synthesizes new viral genomes through a double-stranded RNA (dsRNA) intermediate, which is essential for the further translation and assembly of new virions [4, 5, 7]. In response, the innate immune system detects viral RNA through pattern recognition receptors (PRRs) such as RIG-I and MDA5, which recognize viral RNA features like 5^*′*^-triphosphate ends or double-stranded RNA (dsRNA) intermediates [8, 9] and activate the mitochondrial antiviral signaling protein (MAVS) [10, 11]. The dsRNA intermediate serves as a trigger for the host innate immune system, activating kinases such as TBK1 and IKK*ϵ* via MAVS. These kinases phosphorylate transcription factors like interferon regulatory factors (IRF3/7) and NF*κ*B, which translocate to the nucleus to induce the production of type I interferons (e.g., IFN-*α* and IFN-*β*) [8, 9, 12].

As the virus assembles and releases new virions, the host immune system mounts a broader antiviral response via cytokines like IFNs [8, 10, 11]. Secreted IFNs bind to their receptor (IFNAR) on the surface of the same and neighbouring cells, activating the JAK-STAT pathway. Activated STAT1 and STAT2, together with IRF9, form the ISGF3 complex, which translocates to the nucleus to drive the expression of hundreds of interferon-stimulated genes (ISGs). These ISGs encode a diverse set of antiviral proteins that inhibit various stages of the viral life cycle, including entry, translation, replication, and assembly [9, 12]. Given their crucial role in immune signalling, interferons are also employed therapeutically to enhance immune responses. However, extended or recurrent exposure to IFN can lead to reduced cell responsiveness a phenomenon commonly referred to as IFN desensitisation. The diminished efficacy of IFN’s antiviral function is largely attributed to negative regulators acting at the IFN receptor level [9, 13].

The complex interactions between viral mechanisms and host immune defenses result in dynamic and often unpredictable infection outcomes. The reciprocal actions between the virus and immune host cells establish intricate feedback loops, producing duality in the infection trajectories. For instance, ISGs such as Mx (myxovirus resistance) and IFITM proteins inhibit viral entry, while ISG20, 2’,5’-oligoadenylate synthetase (OAS)-regulated ribonuclease L (RNaseL), IFIT1, and viperin facilitate RNA degradation, inhibit translation, and disrupt egress steps [9, 12]. On the other hand, (+)RNA viruses have also evolved multiple immune evasion strategies to counteract host defenses. These include direct cleavage or degradation of key signaling proteins such as MAVS, TBK1, or STATs by viral proteases, interference with PRR activation or localization, inhibition of ISGF3 formation, and modulation of host gene expression to limit ISG output [11, 14, 15, 16]. In addition, minor alterations in the cellular environments or virus–host interactions can influence the balance between viral proliferation and host immune responses. A similar trade-off between the host and the virus has been suggested to explain the markedly different clinical outcomes observed in individuals with minor variations in their immune responses to the same virus [17, 18]. Therefore, understanding the molecular-level interactions that shape infections is essential [19, 20]. Leveraging the knowledge of critical stages in feedback networks that define host-virus interactions can help us devise therapeutic interventions with improved efficacy.

Given the complexity of these networks, mathematical models trying to capture the underlying behavior have focused on specific aspects of host-virus interactions with varying levels of detail. Several of these models on the viral life cycle [21, 22, 23, 24] and the immune signaling pathways [25, 26, 13, 27] reproduce experimental observations and recapitulate the dynamics of viral and cellular processes accurately. For example, Dahari et al. [22] predicted replicase binding bias for the double-stranded RNA intermediate for the HCV life cycle, whereas more recent virus life cycle models [23, 21, 24] have identified bottlenecks that can be exploited for antiviral treatments. Similarly, mathematical models of innate immune signaling have elucidated key features of the JAK-STAT and RIG-I pathways that modulate IFN responses and demonstrate signatures of IFN desensitisation [13, 25, 26]. However, most current models often isolate the viral life cycle or immune response, and the development of models that capture the dynamic interplay of host and viral factors in immune-competent cells has been challenging. While there is growing interest in understanding the viral dynamics within the context of innate immune signaling, most studies employ a simplified and coarse-grained approach [28, 29, 30, 31, 32, 33, 34]. Alternative systems biology approaches rely on protein–protein interaction (PPI) networks to reconstruct molecular interactions and gene regulatory circuits. By mapping the regulatory interactions, signaling PRRs, downstream effectors, and interferon-stimulated genes (ISGs), key regulatory hubs and the broad architecture of innate immune signalling during viral infection have been elucidated [35, 36, 37]. However, these approaches often treat viral life cycle or innate immune signalling as a static or external parameter, rather than as a dynamic, co-evolving process. Consequently, the disruptive action of viral factors on specific immune nodes, such as cleavage of PRR or their adaptor proteins [16, 14] or ISG degradation [38], is overlooked.

In this study, we present a mechanistic model that integrates detailed mathematical representations of the (+)RNA viral life cycle and host innate immune signalling pathways, along with their dynamic coupling via specific molecular-level interactions. Here, we build on our previous generalized (+)RNA virus model that can accurately recapitulate the dynamics across different (+)RNA virus families [24]. We incorporate a comprehensive immune response network, which includes the key components of the RIG-I/IFN axis [39], enabling us to include the inductor and effector functions of the immune network explicitly. We show that cells respond to IFN in a dose-dependent manner and undergo desensitisation under excessive exposure driven by the TYK and IFN receptors. Notably, viral life cycle dynamics are attenuated by the innate immune response, and virus counteractions on the host cell can negate this control. Similarly, distinguishing characteristics of the virus life cycle can determine the impact of the immune response in controlling virus growth. Clustering host factors with similar temporal expression profiles suggests a stage-wise progression of infection through cell state intermediates. Our integrated model also demonstrates clear bifurcation in infection outcomes, a phenomenon that we hypothesize could be an underlying cause of cellular heterogeneity in viral infections [40, 41, 42, 43, 44, 29]. Counteracting effects defined by the efficiency of virus-induced immune suppression and the strength of antiviral activity of the innate immune system determine this ‘all-or-none’ infection behavior. We identify that the kinetics of viral replication and translation, along with the expression of ISG, are key modulators of the transition between these two completely opposite infection states. Building on our understanding of the IFN dynamics, we propose the prophylactic potential of IFN in suppressing infection and demonstrate how co-targeting viral replication or immune suppression mechanisms can enhance its efficacy. Overall, this work underscores how quantitative systems-level modeling can assist in dissecting virus–host dynamics and offer predictive insights for antiviral strategy.

## Results

### A Cellular Model of Virus-Host Innate Immune Interactions

We present a comprehensive mathematical model that provides an integrated framework for studying virus-innate immune interactions at the single-cell level. Building on two foundational models that individually describe the viral dynamics and immune response, [24, 39] respectively, our model describes the combined dynamics of the virus and host immune response in an infected immune cell (Figure 1a). Previously, we had developed a detailed mathematical representation of (+)RNA virus life cycle that describes the intracellular dynamics of (+)RNA viral constituents, including the effects of vesicular compartment formation, viral mutations, and host cell variability [24]. Although this original viral dynamics model provided a quantitative framework for examining viral replication and growth, it lacked the cellular innate immune response. To address this, we have now incorporated the major immune cascade dynamics (inspired from Burkart *et al*. [39]), which models the RIG-I mediated detection of viral dsRNA and downstream signaling, and the JAK-STAT pathways.

**Figure 1.**
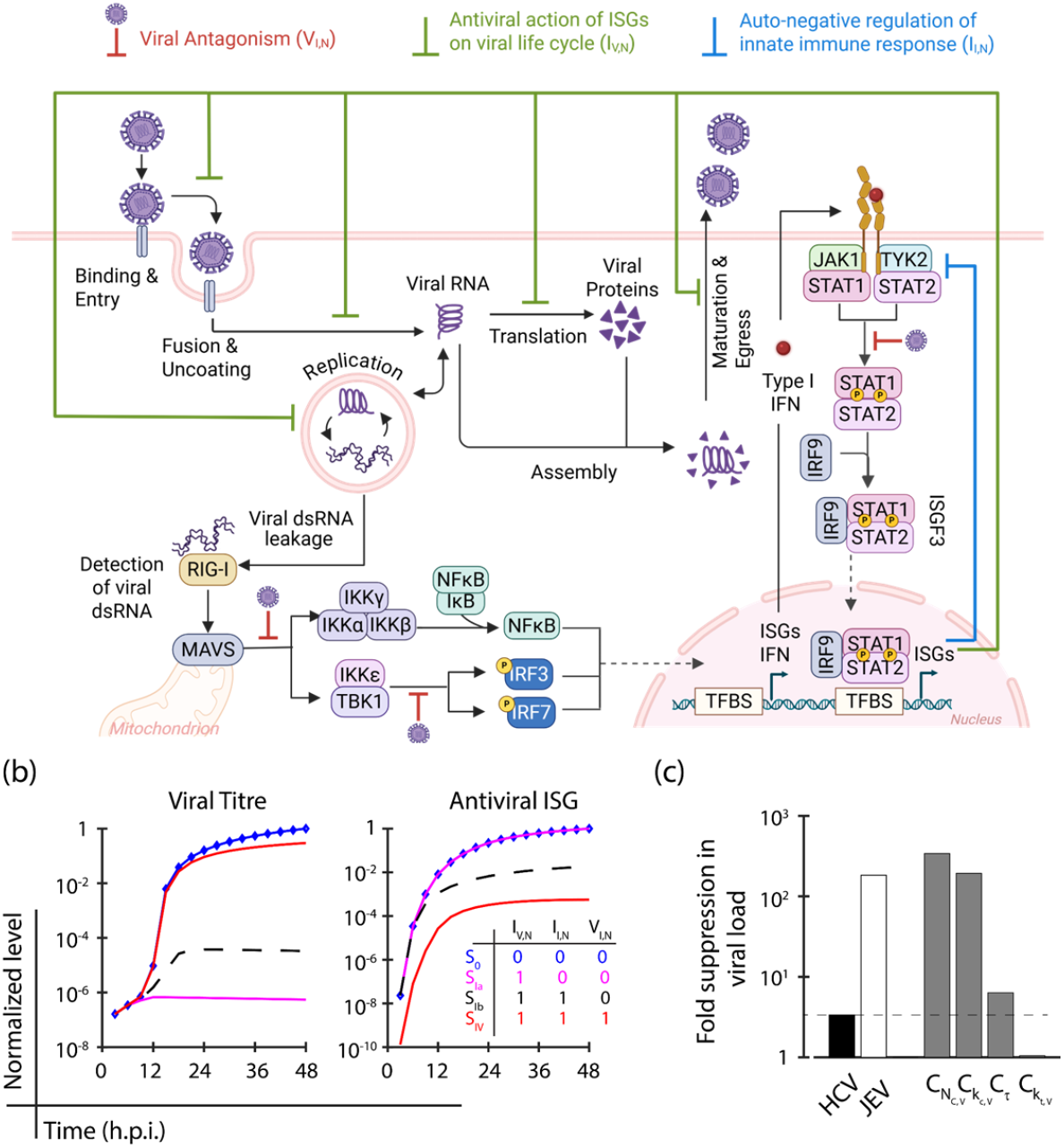
Integrated mathematical model of the viral life cycle with innate immune pathways. (a) Schematic representation of the intracellular model coupling the (+)RNA virus life cycle with host innate immune signalling. Viral infection begins with the extracellular virions (*V*_0_) binding and entering the host cell, followed by uncoating and release of the viral RNA. This RNA undergoes translation to produce structural (*P*_*S*_) and non-structural proteins (*P*_*NS*_), and replication via a double-stranded RNA intermediate (*RC*_*CM*_ ), often sequestered within membrane compartments to evade immune detection. Leakage of replication intermediates into the cytoplasm can trigger recognition by pattern recognition receptors like RIG-I, activating the MAVS and downstream transcription factors such as IRF3/7 and NF*κ*B. This results in the production and extracellular release of type I interferons. Extracellular IFN-I binds to its receptor, activating the JAK-STAT signaling cascade via STAT phosphorylation, which induces the transcription of interferon-stimulated genes (ISGs). ISGs can either suppress the viral life cycle (*ISGav*, strength given by *I*_*V,N*_ ; green repression T-shaped arrows) or auto-regulate the innate immune pathway (*ISGn*, strength parameterized by *I*_*I,N*_ ; blue repression T-shaped arrows). Viral antagonism, particularly via non-structural proteins, can suppress the activity of antiviral ISGs (suppression strength given by *V*_*I,N*_ ; red repression T-shaped arrows). Created with BioRender.com. (b) Viral (left) and ISG (right) dynamics under the influence of various virus-immune interactions, normalized to steady-state levels without virus-immune interactions (*S*_0_: *I*_*V,N*_ = *I*_*I,N*_ = *V*_*I,N*_ = 0, blue). The red line depicts dynamics under the action of viral antagonism and immune suppression along with negative auto-regulation (*S*_*IV*_ : *I*_*V,N*_ = *I*_*I,N*_ = *V*_*I,N*_ = 1). Dynamics with immune activation with antiviral response alone (*S*_*Ia*_: *I*_*V,N*_ = 1, *I*_*I,N*_ = *V*_*I,N*_ = 0, magenta) and host immune activation with negative autoregulation but no virus mediated immune suppression (*S*_*Ib*_: *I*_*V,N*_ = *I*_*I,N*_ = 1 and *V*_*I,N*_ = 0, black) is shown for comparison. (c) Comparison of fold reduction in total viral load (*V*_*T*_ ) between the base case (*S*_0_) and various viruses with the entire virus-immune interaction network (*S*_*IV*_ ) is shown; HCV (black bar), JEV (white bar), and a few hypothetical viruses (grey bars). All the model parameter values of the hypothetical virus, *C*_*θ*_, are based on HCV, except for *θ*, which is representative of JEV. *θ* virus variants were limited to *N*_*C,V*_, *k*_*c,V*_, *τ* and *k*_*t,V*_ since HCV and JEV primarily differ in these parameters.

The integration of viral and immune dynamics was achieved by identifying the mechanisms that links them (a) through viral dsRNA detection that facilitates RIG-I activation of the innate immune cascade, and (b) by detailing the reciprocal interactions between immune response effectors and the viral elements (Figure 1a). During infection, viral double-stranded RNA (dsRNA, the replication intermediate) accumulates in the cytoplasm as it leaks from the replication compartments [45, 46, 15] (equation 4). This cytoplasmic dsRNA becomes accessible to RIG-I, enabling its detection and subsequent activation of the innate immune response. In parallel, ISGs mount a robust and multifaceted defense by targeting viral entry, replication, translation, assembly, and release [9, 12, 47]. To account for these immune responses, we employ a Hill-function-based repression model to represent the ISG-mediated inhibition of these viral processes (Figure 1a, equations 1, 2, 3, 4 and S2- equations, S5, S6, and S7). Apart from the effect of innate immune response on viral dynamics, we also account for how viral proteins actively interfere with the innate immune response by targeting key components of the signaling pathways. To model this interference, we introduce viral protein-mediated repression of signal propagation via MAVS, IKK*ϵ*-TBK1, and the STAT1/STAT2 receptor complexes [48, 49, 50] (Figure 1a, SI equations- S12, S18, S40, S50, and S67). Finally, we incorporate the regulation of IRF7 (equations 5 and 6), a master transcriptional regulator of type I interferons (esp. IFN*α*) previously overlooked in earlier models. IRF7 plays a critical role in amplifying IFN*α* responses through a positive feedback loop mediated by JAK-STAT signaling, where initial IFN*β* production (mediated by IRF3) induces IRF7 expression (details provided in Methods). To precisely capture the dynamics of the cellular immune response, innate immune parameters are adapted from prior studies that estimate them using experiments with synthetic 5’ppp-dsRNA or IFN to stimulate cells [13, 39]. Virus-specific parameters for HCV and JEV are obtained from our earlier work that reconciles experimental data of viral protein and RNA dynamics with viral life cycle dynamics [24].

To investigate the broad impact of the bidirectional virus-host interactions on infection progression, we first employ an aggregated parameter approach to examine the viral-immune dynamics in our comprehensive integrated model. For this purpose, we introduced two lumped tuning parameters: *I*_*V,N*_, representing the strength of immune-mediated suppression of the viral processes (equations 1, 2, 3, 4 and SI equations- S2, S5, S6, and S7; see Methods), and *V*_*I,N*_, capturing the extent of viral protein-mediated suppression of the immune response (SI equations- S12, S18, S40, S50, and S67; see Methods). Additionally, *I*_*I,N*_ serves as a tuning parameter to modulate the negative feedback effect of anti-inflammatory ISGs on the JAK-STAT pathway (SI equations- S32, S39, S40, S42, S43, and S44; see Methods). At the molecular level, these interactions influence different processes such as immune sensor activation, signaling, viral replication, and protein translation to varying extents; however, for analytical simplicity, we will assume a uniform regulatory effect throughout for each of the immune-mediated suppression and the viral antagonism processes. This approach ensures that the model remains interpretable while allowing us to explore the broader implications of these interactions on the infection outcomes. We begin by defining a baseline scenario (*S*_0_) devoid of counteracting interactions between the immune or viral components (*I*_*V,N*_ = *V*_*I,N*_ = *I*_*I,N*_ = 0, Figure 1b). Under this condition, the model successfully reproduces the dynamics of Hepatitis C virus (HCV) proteins, consistent with experimental observations [23, 24]. In this scenario, immune pathways are gradually activated, as evidenced by the increase in levels of interferon- stimulated genes (ISG). Next, we consider the scenario where the antiviral effects of the innate immune response are included (*S*_*Ia*_: *I*_*V,N*_ = 1, *I*_*I,N*_ = *V*_*I,N*_ = 0). Here, while innate immune dynamics remain unchanged, the viral infection is significantly disrupted. In the subsequent scenario, we incorporate the negative self-regulation of the innate immune pathway (*S*_*Ib*_: *I*_*V,N*_ = *I*_*I,N*_ = 1; *V*_*I,N*_ = 0). This results in a dampened immune response, leading to a less pronounced reduction in viral load compared to *S*_*Ia*_. Finally, we examine the complete system by including the virus-mediated suppression of the innate immune pathways with the combined action of the immune pathways (*S*_*IV*_ : *I*_*V,N*_ = *I*_*I,N*_ = *V*_*I,N*_ = 1). In this scenario, the immune response is further suppressed, as indicated by a significant reduction in ISG levels. In fact, steady-state ISG expression in *S*_*IV*_ is approximately four orders of magnitude lower than in the baseline scenario (*S*_0_), suggesting strong repression of the immune response due to virus interference. However, there is only a modest change in the level of virus growth across the two scenarios (Figure 1b). This shows how potent viral suppression mechanisms can significantly attenuate the immune response while maintaining viral persistence.

By adjusting the model parameter values (SI table S4) to reflect the dynamics of Japanese Encephalitis Virus (JEV) [24], the system predicts kinetics that are qualitatively similar to those observed for HCV (SI Figure S1). However, virus-specific differences emerge when comparing the steady-state values across different scenarios. The immune-mediated suppression of viral infection as indicated by the steadystate viral load ratio *S*_0_*/S*_*IV*_, is markedly higher for JEV (182.6) compared to HCV (3.4), highlighting a stronger suppression effect on JEV than HCV (Figure 1c). We hypothesize that this disparity arises from key differences in the viral life cycle parameters of JEV. Notably, JEV exhibits a faster formation rate of replication complexes (higher *k*_*c,V*_ ) and larger number of replication complexes (higher *N*_*C,V*_ ) compared to HCV. These factors drive a more rapid and substantial accumulation of replication intermediates, which in turn facilitates earlier and stronger activation of the immune response. Consequently, when the antiviral effects of the immune response are incorporated (*S*_*IV*_ ), the reduction in viral load is significantly greater for JEV than for HCV. This highlights the critical role of viral replication dynamics in shaping the interplay between viral persistence and immune suppression.

To validate this hypothesis, we constructed a panel of hypothetical viral strains, each retaining the characteristics of HCV except for a single substituted feature that is derived from JEV (Figure 1c). For example, the strain *C*_*τ*_ that exhibits JEV-like vesicular compartment formation timescale showed a modest (*<*10-fold) suppression of viral load in scenario (*S*_*IV*_ ) compared to the baseline scenario (*S*_0_). However, the strains *C*_*NC,V*_ and *C*_*kc,V*_ which exhibit JEV-like carrying capacity (*N*_*C,V*_ ) and formation kinetics of replication complexes (*k*_*c,V*_ ), respectively, show over 200-fold suppression of viral load. Therefore, the HCV chimeric strains with higher *N*_*C,V*_ and a larger *k*_*c,V*_ are similar to JEV rather than wild-type (wt)HCV, suggesting that these two viral growth characteristics of JEV are responsible for its enhanced suppression by the innate immune response. These results support our hypothesis that early and robust immune activation, driven by faster and larger accumulation of viral dsRNA in the cytoplasm, leads to more effective suppression of the viral life cycle, consistent with observations in other systems [51]. In contrast, a HCV chimera with JEV-like translation rate (*C*_*kt,V*_ ; 3-fold faster translation compared to wt-HCV) exhibited negligible difference in viral suppression. The diminished suppression ratio of *C*_*kt,V*_ probably results from an early and increased synthesis of viral proteins, allowing for quicker and more effective inhibition of host immune components.

### Reciprocal Negative Feedback between Viral and Immune Processes Shapes the Viral-Immune Bifurcation

Experimental studies have demonstrated that early and robust innate immune responses, particularly those mediated by interferon-stimulated genes (ISGs), are essential for limiting viral replication and achieving viral clearance [52, 53]. In contrast, delayed or dysregulated immune responses, often resulting from viral immune evasion strategies, can lead to immune escape and prolonged infection [54, 55, 56, 57]. Both the immune suppression exerted by viruses and the strength of host innate immune responses are known to vary continuously, influenced by factors such as viral load, replication kinetics, host cell type, and genetic background. For example, different viral strains can differ in their ability to produce immune antagonists or stimulate ISG expression, while host immune activity is modulated by polymorphisms in interferon signaling components, ISG expression profiles, and metabolic or stress conditions. Motivated by these biological complexities, we investigated whether our virus-immune model is capable of capturing the dichotomy in infection outcomes and identifying the key cellular and viral features that govern the balance between viral clearance and immune escape.

The strength of viral antagonism (*V*_*I,N*_ ) determines distinct immune response dynamics. At low *V*_*I,N*_ values, ISG levels remain high and the viral loads remain low, indicating robust immune activation and effective viral clearance. However, as *V*_*I,N*_ increases, ISG levels are progressively suppressed, particularly at higher viral loads, reflecting the dominance of viral antagonism and the transition to immune escape (Figure 2a). For HCV, the infection outcome switches from a ‘virus-low, ISG-high’ state to a ‘virus-high, ISG-low’ state (Figure 2a) as *V*_*I,N*_ surpasses 0.44, suggesting that HCV can overcome immune control above this critical level. Further, a threshold viral load is evident below which ISG levels remain low, regardless of *V*_*I,N*_, suggesting that such low viral loads are insufficient to trigger strong immune activation.

**Figure 2.**
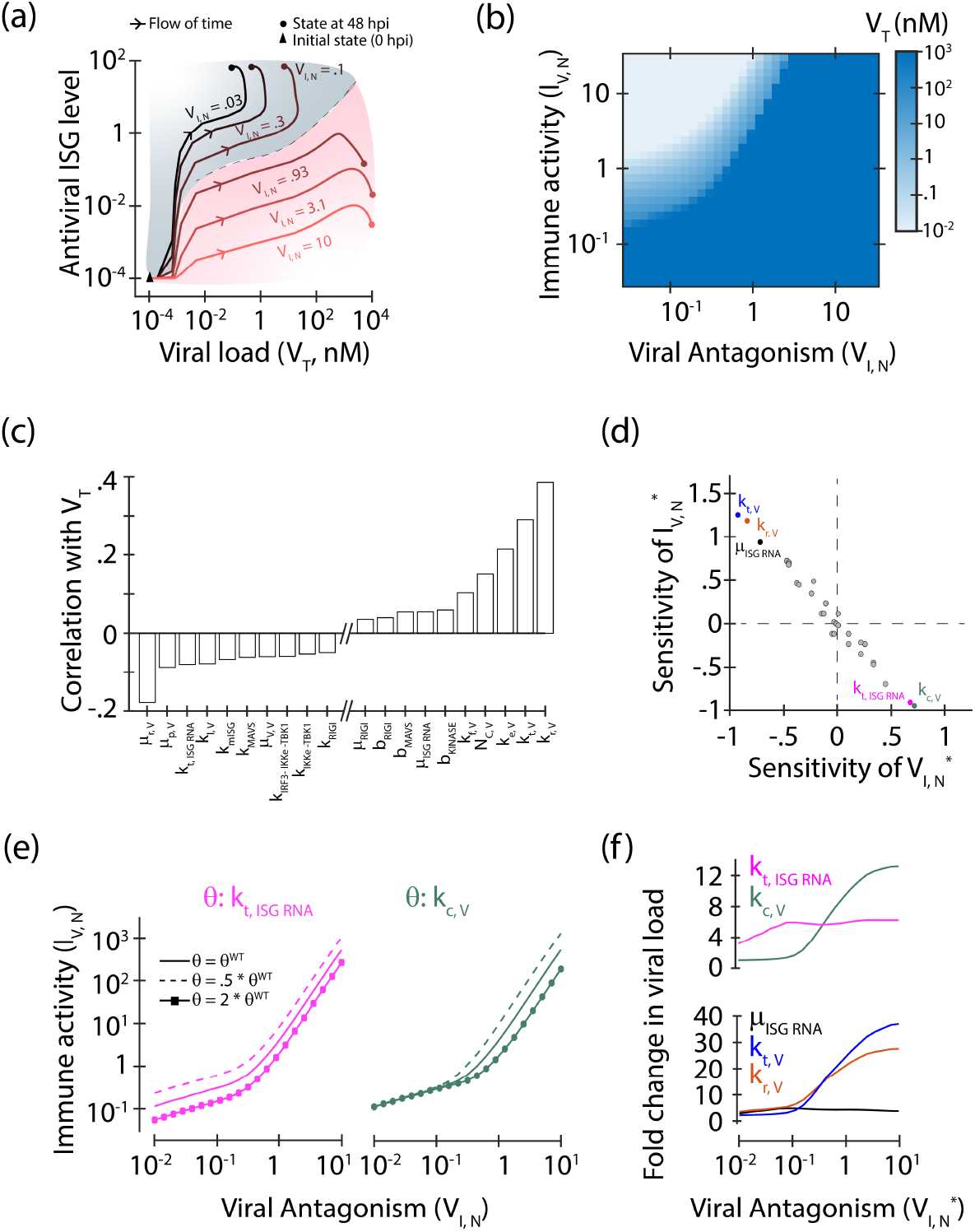
Virus-immune bifurcation and sensitivity analysis reveal critical regimes in infection. (a) The phase plot showing the dynamics of viral load and antiviral ISG levels during hepatitis C virus infection is shown for various strengths of viral antagonism (*V*_*I,N*_ ). The system shifts from a ‘virus-high, ISG-low’ state (pink region) to a ‘virus-low, ISG-high’ state (grey region) as *V*_*I,N*_ increases, with the clear transition at 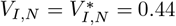. (b) The viral load (*V*_*T*_ at 48 h.p.i) HCV is shown as a function of *V*_*I,N*_ and *I*_*V,N*_ . (c) Correlation coefficients between viral load (*V*_*T*_ ) and individual parameters near the *V IB* boundary shows how a parameter influences the steady-state viral load, thus identifying antiviral and proviral parameters. (d) Parameter sensitivity indices for 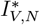 and 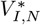 evaluated at the bifurcation point: 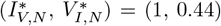 are negatively correlated. Colored dots denote model parameters with threshold sensitivity score (see Methods) *>* 1. (e) The *V IB* boundary for HCV infection systems is shown for different values of *k*_*t,ISGRNA*_ (left) and *k*_*c,V*_ (right). The solid line, dashed line, and the solid line with square symbols show the *V IB* boundary when the parameter is unperturbed, reduced by half, and doubled, respectively. (f) Fold change in the steady state HCV viral load when the corresponding parameter values were (top panel) halved (magenta-*k*_*t,ISGRNA*_, green-*k*_*c,V*_ ), or (bottom panel) doubled (black-*µ*_*ISGRNA*_, blue-*k*_*t,V*_, and orange-*k*_*r,V*_ ) along the *V IB* boundary.

We can now evaluate how virus infection is established as a function of the strength of the two key regulatory types of interactions in our model: the immune-mediated viral suppression (by varying *I*_*V,N*_ ) and the viral antagonism of the immune response (by varying *V*_*I,N*_ ). The steady-state viral load (*V*_*T*_ ) as a function of immune activity (*I*_*V,N*_ ) and viral antagonism (*V*_*I,N*_ ) reveals a bifurcation boundary separating two distinct regimes: (a) a low-virus regime, where high *I*_*V,N*_ and low *V*_*I,N*_ result in effective immune clearance, and (b) a high-virus regime, where low *I*_*V,N*_ and high *V*_*I,N*_ lead to immune escape (Figure 2b). The set of critical points 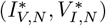 where such transitions occurs in the *I*_*V,N*_ *− V*_*I,N*_ parameter space, captures the tipping point between innate immune control and virus escape and is referred here as the Viral-Immune Bifurcation (*V IB*) boundary. There is a marked positive correlation between the critical values of 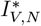 (immune activity) and 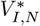 (viral antagonism), indicating a balancing interaction: as viral antagonism intensifies, enhanced immune activity is necessary to counteract it. Overall, the *V IB* boundary emphasizes the nonlinear dynamics of the virus-immune system, where robust immune activation (high ISG levels) can suppress viral replication, but strong viral antagonism can overwhelm the immune response, leading to persistent infection.

An analogous trend is detected for JEV (SI Figure S3). However, with the same level of innate immune activity (*I*_*V,N*_ = 1), JEV requires a lower level of antagonism to evade the immune activity 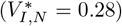 compared to HCV 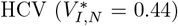. The remarkable capacity of JEV to overcome immune clearance, despite significant suppression by the host immune response (Figure 1c), can be attributed to its substantially higher rate of protein synthesis. This rapid translation expedites the availability of viral proteins to amplify the viral antagonism, enhancing immune evasion. In contrast, HCV’s slower protein synthesis [24] demands a more robust and efficient antagonistic strategy to achieve comparable immune suppression.

Our initial approach involved broad perturbations across multiple interactions (varying *I*_*V,N*_ and *V*_*I,N*_ ) rather than specific molecular-level processes. However, such simultaneous perturbations are unlikely to occur in natural systems. A more plausible scenario is that the fitness of the virus or the immune response is influenced by specific changes in a single or a small subset of parameters. To explore this, we investigated whether the Virus-Immune Boundary (*V IB*) emerges naturally from the virus-immune interaction network under more realistic conditions, where only a limited set of parameters change for different viral strains or cell types. Notably, similar sharp transitions between viral clearance and escape states with distinct *V IB* boundaries were observed for HCV when we varied specific model parameters, such as *k*_*t,V*_, *k*_*c,V*_ and *k*_*ISG RNA*_ (SI Figure S2). This sharp transition at the *V IB* boundary provides a unique opportunity to dissect the relative contributions of viral and host processes to infection outcomes without the need to simulate the full system dynamics under individual parameter variations.

By leveraging the system’s heightened sensitivity near the *V IB* boundary, we can identify key parameters that strongly influence infection outcome. The analysis in Figure 2(c) illustrates how this approach enables the identification of critical drivers of the bifurcation (at 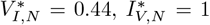 for HCV). Specifically, we simulate the infection outcomes using a set of 25,000 parameter combinations, where each parameter is sampled from a *±*10% range of their nominal values (see SI Table S3, S5-S8, and Methods). This approach allows us to recover a bimodal distribution in viral loads (SI Figure S4) that highlights the pronounced dichotomy in infection states near the *V IB* boundary. Approximately 15% of the parameter combinations lead to viral clearance, while a large fraction of parameter sets result in high viral load (*>* 10^2^ nM), suggesting immune escape. Correlation between the sampled parameter values and the resulting viral load helps us in identifying proviral and antiviral factors (Figure 2c). We find that the degradation rate of viral RNA (*µ*_*r,V*_ ), proteins (*µ*_*p,V*_ ) or extracellular virus particles (*µ*_*V,V*_ ) is strongly correlated with reduced viral load, highlighting their obvious role as critical parameters controlling viral growth. Similarly, parameters that enhance viral processes—such as viral translation (*k*_*t,V*_ ), replication (*k*_*r,V*_ ), export of viral (+)RNA from replication compartments (*k*_*e,V*_ ) or the carrying capacity of compartmentalized replication complexes (*N*_*C,V*_ ) - are positively correlated with higher viral loads. On the other hand, enhancing immune detection, for example, via increasing the leakage of compartmentalized viral dsRNA into the cytoplasm (*k*_*l,V*_ ), leads to a reduction in viral load. Among the host cell properties, potent immune signaling and activation can similarly reduce virus production. For instance, rapid activation of IRFs (*k*_*IRF* 3*−IKKe−T BK*1_), enhanced phosphorylation of TBK1 or IKK*ϵ* (*k*_*IKKe−T BK*1_), or increased activation rates of MAVS or RIG-I (*k*_*MAV S*_, *k*_*RIGI*_ ) or upregulation of host ISG mRNA and protein (*k*_*mISG*_, *k*_*t,ISGRNA*_) can considerably decrease viral load. Conversely, when immune factors undergo rapid removal due to enhanced degradation or deactivation, we observe increase in viral load suggesting diminished innate immunity. This is exemplified for the case of RIG-I (*µ*_*RIGI*_ ) or ISG mRNA (*µ*_*ISGRNA*_) degradation, and dephosphorylation of RIG-I (*b*_*RIGI*_ ), MAVS (*b*_*MAV S*_), or kinases (*b*_*KINASE*_).

To further dissect the mechanisms governing the balance between immune and viral interplay, we performed a bifurcation sensitivity analysis to quantify how changes in crucial viral and host parameters affect the system’s tipping point (details in Methods). A distinct inverse correlation is evident in how various parameters affect 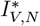 and 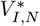, underscoring the antagonistic nature of these two sets of interactions (Figure 2d). Interestingly, some molecular-level processes stand out as more proviral or antiviral than the others. Among the viral processes, parameters that directly influence protein production, such as translation rate *k*_*t,V*_, or indirectly by boosting RNA synthesis, like replication rate *k*_*r,V*_, exhibit the strongest negative influence on 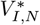 (SI Figure S5). For instance in the case of HCV, a +10% increase in *k*_*r,V*_, reduces the viral antagonism 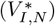 required for virus escape by 6.8% (from 0.44 to 0.41) for a fixed 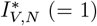. Thus, enhanced viral replication and translation aid in efficient immune escape. Regarding the immune response, mechanisms that enhance ISG activity (*µ*_*ISGRNA*_ and *k*_*t,ISGRNA*_) reduce the required strength of the immune response to control the viral proliferation. Conversely, we observe that enhanced viral replication compartmentalization can decrease 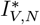. This paradoxical antiviral nature of replication compartmentalization in assisting the immune response can be attributed to the concomitant increase in leaked cytoplasmic dsRNA, leading to a prominent immune activation. For instance, at the *V IB* boundary, when the rate of compartmentalized replication complex formation (*k*_*c,V*_ ) is increased by +10%, the antiviral activity 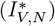 required for immune control is decreased by 6% (from 1 to 0.94) for HCV. This provides additional evidence that elevated *k*_*c,V*_ may trigger earlier and more abundant production of immune agonists (dsRNA), which support the immune response (Figure 2c). In case of JEV, swift formation of replication compartments (small *τ* ) result in rapid dsRNA accumulation. This triggers a potent immune activation and requiring only mild immune response to suppress it. Conversely, in HCV infection, slower dsRNA production postpones immune activation, necessitating stronger immune suppression to control infection.

We next extend our bifurcation sensitivity analysis to investigate such virus-immune tipping point dependence on various critical cellular and viral processes (identified in Figure 2d) along the entire *V IB* boundary. Figure 2e (and SI Figure S6 and S7) illustrates how a twofold increase or a 50% decrease in ISG translation efficiency or viral replication complex formation influences the *V IB* boundary across the entire range of viral antagonism 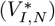 and immune activity 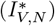 for HCV. Higher ISG RNA translation rate (*k*_*t,ISGRNA*_) requires lower immune activity to counteract viral antagonism (downward shift in *V IB* boundary) and achieving viral clearance, while reducing it leads to weakening of immune control (upward shift in *V IB* boundary) (Figure 2e left). We observe that a two-fold increase in ISG translation (*k*_*t,ISGRNA*_) leads to a 3-6 fold decrease in HCV viral load at different points on the bifurcation boundary (Figure 2f top). Effectively, an increase in *k*_*t,ISGRNA*_ leads to a downward shift of the *V IB* boundary in the *I*_*V,N*_ *− V*_*I,N*_ space, indicating that change in the host ISG RNA translation rate consistently impact the system’s behavior uniformly along the bifurcation boundary. This trend is also observed for a two-fold higher ISG mRNA degradation rate that results in an *∼*3-4 fold increase in viral load at different points along the *V IB* boundary.

Unlike host-mediated processes, perturbations in viral processes, such as replication (*k*_*r,V*_ ), translation (*k*_*t,V*_ ), or replication compartmentalization (*k*_*c,V*_ ), influence system behavior in a regime-dependent manner across the bifurcation interface (Figure 2e, right; SI Figure S6). At high levels of both viral antagonism and immune activity, increasing the rate of replication compartmentalization (*k*_*c,V*_ ) shifts the *V IB* boundary upward, requiring stronger immune activity (*I*_*V,N*_ ) to counteract viral antagonism (*V*_*I,N*_ ) and reduced levels of viral replication complex compartments makes it easier for the immune system to suppress viral replication. Similar asymmetric changes in the *V IB* are also observed when other virus-mediated proviral processes are impeded (SI Figure S7). Such dependence of these viral processes even influences the viral load levels in a regime-dependent manner along the *V IB* boundary (Figure 2f). When virus–immune interactions are weak (low *I*_*V,N*_ and *V*_*I,N*_ ), viral load (*V*_*T*_ ) is largely insensitive to changes in *k*_*c,V*_, *k*_*r,V*_, or *k*_*t,V*_ (Figure 2f). However, in the strongly interacting regime of the *V IB* boundary, small changes in the proviral factors have disproportionately large effects. For instance, along the boundary at high *V*_*I,N*_ (= 10), a two-fold decrease in *k*_*c,V*_ results in a *∼*12-fold increase in *V*_*T*_, while a two-fold increase in *k*_*r,V*_ or *k*_*t,V*_ yields *∼*30-fold and *∼*40-fold increases in *V*_*T*_, respectively (Figure 2f). Comparable trends for JEV (SI Figures S6, S7 and S8) suggest this regime-dependent sensitivity is a general feature of (+)RNA virus-host immune interactions.

These regime-dependent effects of viral parameters can be explained by two factors: (a) the dual role of viral dsRNA as both a viral replication intermediate and an immune activator, and (b) the double-negative feedback loop between viral antagonism (*V*_*I,N*_ ) and immune suppression of viral replication (*I*_*V,N*_ ). Given the dual role of viral dsRNA, changes in viral processes, like viral replication, translation, and replication compartmentalization, that alter its levels exert both pro-viral and antiviral influences. These counter-influences tend to balance each other at the *V IB* interface. When the negative feedback interaction strengths are high, strong mutual antagonism amplifies even minor imbalances, producing pronounced shifts in infection dynamics and outcome (Figure 2e and f).

### Dynamic Phases of Immune Response during Infection Progression

Deciphering the temporal dynamics of immune responses during infection is essential for identifying key regulatory mechanisms, critical immune checkpoints, and potential therapeutic targets. In complex biological systems like viral infections, immune factors and viral components interact over distinct temporal phases, each defined by specific molecular patterns. However, due to the vast array of immune components, monitoring the dynamics of all elements is impractical in experimental settings, and identifying the most informative markers poses a complex challenge. We can leverage our virus-immune response model to address this challenge.

Here, we investigate the temporal profiles of immune components and examine their temporal correlations to pinpoint effective indicators of cellular infection spread. To capture the immune system’s underlying dynamics, we examine its behavior under three distinct virus-immune regimes (Figure 3a and SI Figures S9-S11): (a) at the *V IB* boundary 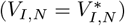, (b) viral clearance regime 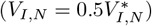, and (c) immune escape regime 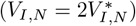. Hierarchical clustering of the z-normalized trajectories of immune factor dynamics across these regimes revealed eight distinct groups of dynamically co-regulated immune factors (Figure 3a). Each cluster exhibits unique temporal patterns that can be delineated into three major temporal phases of immune response during infection: an ‘early activation (EA)’ phase (0–12 h), an ‘intermediate response (IR)’ phase (12–24 h), and a ‘late resolution (LR)’ phase (24–48 h) (Figure 3b).

**Figure 3.**
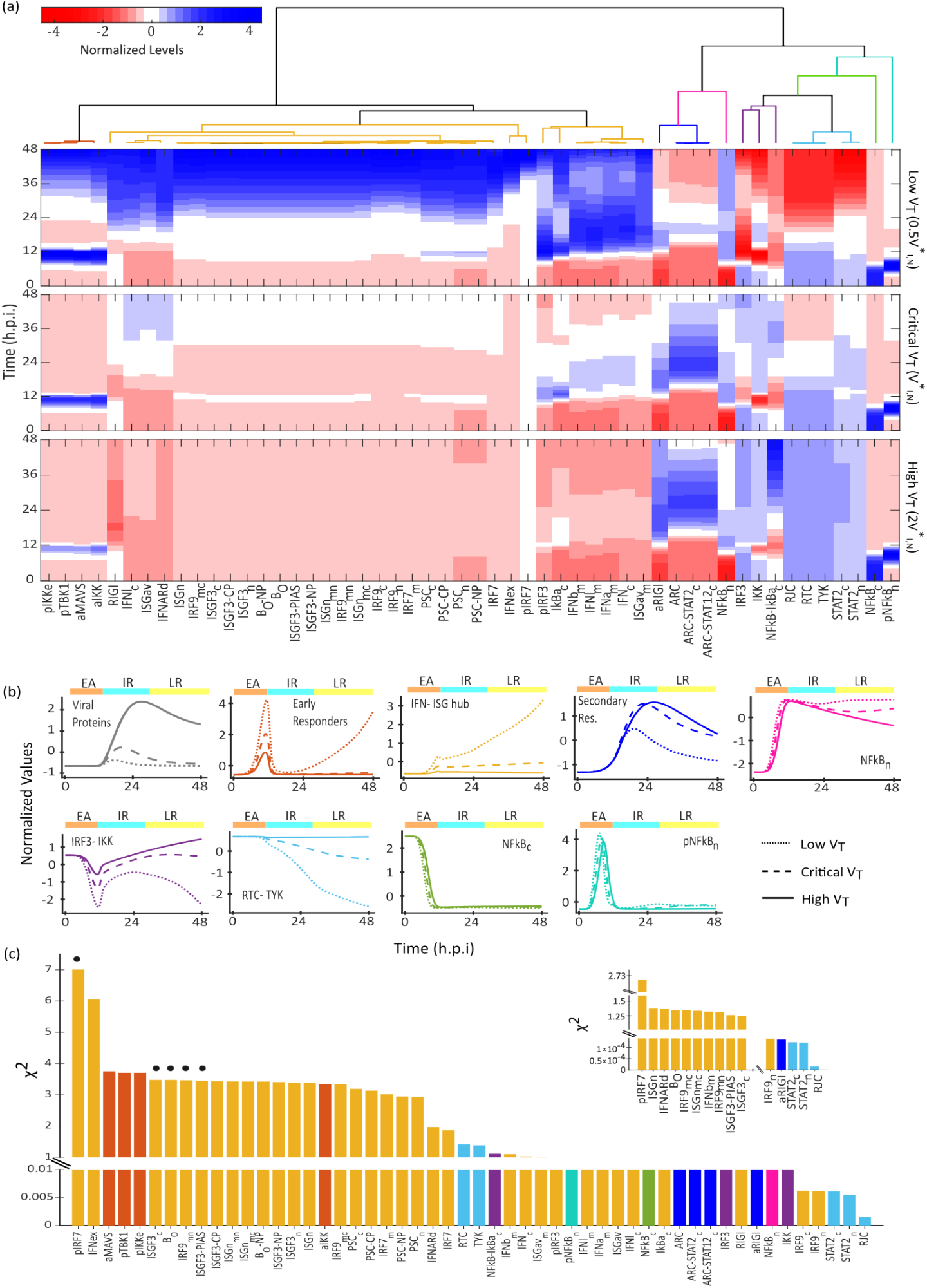
Dynamics of innate immune effectors under different infection outcomes. (a) Clustered z-normalized dynamics of immune pathway components under three viral-immune regimes: low *V*_*T*_ (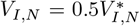; top), critical *V*_*T*_ (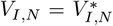; middle) and high *V*_*T*_ (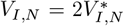; bottom) are plotted. Distinct clusters are indicated by the color-coded dendrogram. (b) The panels show the dynamics of viral proteins (grey) and the mean temporal profiles of the immune pathway clusters identified in (a), revealing their characteristic dynamic behavior under each regime. (c) The *χ*^2^ metric, quantifying the divergence between dynamics of immune factors across the low *V*_*T*_ and the high *V*_*T*_ regimes, is plotted (see Methods; bar edge colors reflect the cluster assignment). The inset shows the *χ*^2^ metric for the early phase (0–16 h). Dots indicate factors that show high divergence in both time periods.

In the ‘early activation’ phase, the newly formed viral dsRNA is detected by the RIG-I, triggering a sudden but transient increase in the levels of the “Early responders” cluster (Figure 3a and b). This group is dominated by the activated forms of antiviral signal transducers (aMAVS, pIKK, pTBK1, pIKKe). Their levels reach a peak around 8 to 12 hours, followed by a sharp decay. Overall, “Early responders” display similar initial transient dynamics in all three infection regimes. However, their induction level is inversely proportional to the severity of the infection, i.e., strong activation of the “Early responders” is associated with the viral clearance regime. Additionally, “Early responders” increase further in the resolution phase, likely due to reinforcement of immune signaling when initial immune responses are effective.

During the ‘intermediate response’ phase, as the viral replication is established and viral dsRNA increases, RIG-I activation increases. This initiates the activation of the “IFN-ISG transcriptional hub” (Figure 3a and 3b), which encompasses phosphorylated STATs, IRFs, and IFN/ISG mRNAs. These immune components exhibit a progressive increase in their levels as the infection advances, reaching their highest concentration during the ‘intermediate response’ phase. However, the increase was substantially repressed under high *V*_*T*_ (immune escape regime), which can be attributed to the suppression of the signaling pathways by the viral proteins (Figure 1a). Conversely, when considering viral clearance characterized by low *V*_*T*_, the elevation in levels during the resolution phase is significantly greater than in other regimes. This suggests robust reinforcement of pathways and effective antiviral transcriptional activity.

In the ‘late resolution’ phase, the IFN receptor together with the JAK-STAT pathway plays a pivotal role in directing immune responses. This activation is triggered through the formation of complexes that include kinases, specifically TYK, along with IFN receptors like RTC and RJC, and STATs. This collective assembly is referred to as the “TYK-RTC” cluster (Figure 3a and b). The behavior of this group of immune modulators exhibited a gradual decrease throughout the infection period, potentially as a result of feedback inhibition and resource depletion. However, the reduction was less evident in high viral load conditions due to diminished upstream engagement caused by the robust immune suppression by the virus. Activation of “TYK-RTC” group leads to activation of “Secondary responders”, comprising of STAT-bound IFN receptor complexes peaking during the late phase. This cluster displayed a delayed yet persistent activation at high *V*_*T*_, possibly suggesting a partial, ineffective triggering of IFN signaling that does not effectively counteract viral suppression. Furthermore, the “IRF3-IKK” cluster, comprising of the native, unphosphorylated forms of key signaling intermediates like IRF3 and IKK, exhibits inverse dynamics, with higher levels observed under high *V*_*T*_ . This is likely due to impaired activation, suggesting ineffective immune progression.

*NFκB* dynamics during virus infection can be understood from the behaviour of the “*NFκB*_*c*_”, “*pNFκB*_*n*_”, and “*NFκB*_*n*_” forms (Figure 3a and b). Cytoplasmic “*NFκB*_*c*_” is quickly depleted due to phosphorylation, nuclear import, and its capture by I*κ*B*α*. This rapid drop highlights its role as the input pool for downstream *NFκB* signaling. As “*NFκB*_*c*_” translocates and phosphorylates into “*pNFκB*_*n*_”, it induces IFNs and feedback regulator I*κ*B*α*. The transient peak of “*pNFκB*_*n*_” is concomitant with “Early responders” group dynamics, peaking around 8-12 hours followed by a sharp decay. Finally, quick dephosphorylation of “*pNFκB*_*n*_” gives rise to “*NFκB*_*n*_”, which rises shortly afterwards, then decays gradually over the course of the infection due to cytoplasmic translocation and I*κ*B*α* feedback. At high *V*_*T*_, it continues to decay gradually, reflecting a partially active and inefficient *NFκB* cycle caused by viral suppression of the MAVS–IKK signaling pathway. However, under low viral load, “*NFκB*_*n*_” levels can gradually increase during the late resolution (LR) phase owing to enhanced signaling activity.

Taken together, our analysis reveals a hierarchical temporal cascade in the host antiviral response, wherein identified clusters reflect functionally distinct stages of innate immune signaling and effector activation. The clusters show significant sensitivity to viral load, with their rapid modulation highlighting crucial points where host detection and response pathways may succeed or fail. Notably, the early divergence of the “IFN-ISG transcriptional hub” in various infection regimes suggests these molecules can serve as early indicators of infection outcomes, even before clear phenotypic signs appear.

We employ a *χ*^2^ metric to compare the difference between high and low viral load scenarios (Figure 3c, see Methods). Phosphorylated IRF7, nascent IFN transcripts, and early ISGs are key factors in viral infection outcomes, particularly in the initial stages of infection, indicating that the immediate-early transcriptional circuit plays a crucial role in initiating effective antiviral states (Figure 3c-inset). Throughout the infection period, the persistent impact of IFN levels, phosphorylated IRF7, and activated signaling kinases like aMAVS and pIKK highlights the critical roles of strong signal initiation and downstream amplification in immune regulation (Figure 3c). These findings map the dynamics of viral antagonism directly onto distinct regulatory layers of the host response, illustrating how viral interference at both early detection and later transcriptional amplification stages can rewire immune trajectories and shape disease outcomes. Our model facilitates the understanding of the hierarchical and temporal structure of the antiviral response, identifying specific molecular clusters as crucial indicators of the infection progression.

### IFN Desensitization is Mediated by Receptor-Kinase Complex and TYK Dynamics

IFNs are widely used as therapeutic agents for the treatment of chronic viral infections, certain cancers, and autoimmune diseases, due to their potent antiviral and immunomodulatory effects [58, 59]. However, their clinical efficacy is often limited by the phenomenon of desensitization, where repeated or prolonged IFN exposure leads to reduced cellular responsiveness [60]. This behavior is thought to arise from feedback inhibition and regulatory dynamics within the JAK-STAT signaling pathway [61, 62, 63].

Because our model incorporates key immune pathways that regulate the IFN response, it can capture the dose-dependent desensitization effect for a naive (uninfected) cell (see Methods). A high dose of IFN (100 nM) sharply induces ISGs responses with ‘ISGn mRNA’, an early transient negative regulator [13], and the ‘ISGav mRNA’, a late antiviral transcript [13], quickly rising to 0.75 nM and 140 nM, respectively (Figure 4a and b). This activation is brief, with both mRNA dropping significantly and stabilizing at approximately 0.01 nM and 60 nM within 24 hours. A second high (100 nM) IFN dose 24 hours later elicits no additional increase in ISGn or ISGav mRNA, signifying strong desensitization. Conversely, administering a high subsequent IFN dose (100 nM) following an initial low dose (1 nM) results in an increase in ISGn and ISGav, albeit it is modest compared to the substantial increase observed with a single high-dose (100 nM) of IFN. This signifies limited IFN desensitization consistent with experimental reports of dose- and time-dependent IFN desensitization [27, 13].

**Figure 4.**
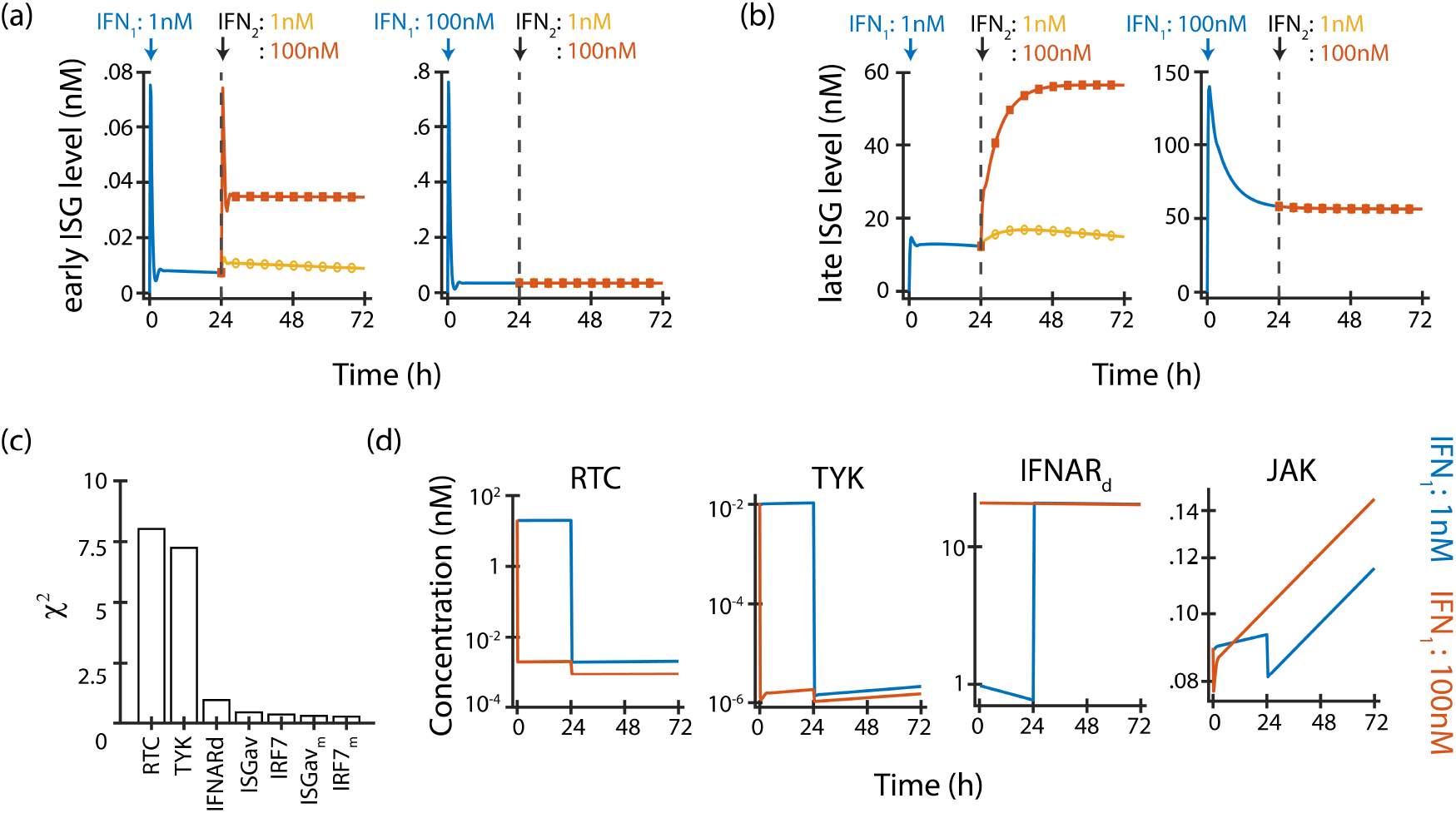
Dose-dependent IFN-desensitization. The kinetics of (a) an ISG transcript involved in immune suppression (early ISG, ISGn) and (b) an ISG transcript involved in viral suppression (late ISG, ISGav) are compared under distinct two-dose interferon (IFN) treatments. The second dose is administered 24 hours after the initial dose. The left panels represent Condition 1, where an initial low dose of 1 nM IFN is followed by either a low (1 nM, yellow line) or high (100 nM, orange line) second dose.The right panels represent Condition 2, where an initial high dose of 100 nM IFN is followed by either a low (1 nM, yellow line) or high (100 nM, orange line) second dose. (c) The *χ*^2^ values, representing deviations in the dynamics of JAK-STAT pathway components between the two IFN priming conditions, are plotted. The top-ranking species, which indicate model components most sensitive to the initial IFN dose, are highlighted. (d) Time- course dynamics of select model species with the two highest (*χ*^2^: RTC, TYK) and two moderate (*χ*^2^: IFNARd, JAK) values are shown, illustrating IFN dose-dependent differences in system response.

Comparing these scenarios, most downstream signaling components remain similar at re-exposure, but key upstream regulators, particularly TYK2 and the IFN receptor complex (IFNAR1/2), show marked depletion after high-dose exposure (Figure 4c, d). This indicates that desensitization arises from the destabilization or inhibition of receptor-proximal components, as suggested by experimental reports [13, 62, 61, 63]. Our earlier analysis also grouped TYK2 and the receptor complex within the same regulatory set (Figure 3), reinforcing their central role in signal initiation and their vulnerability to high-dose IFN exposure.

To further test this hypothesis, we simulated a two-dose IFN regimen (100 nM followed by 100 nM) and selectively restored the RTC, TYK, and JAK levels prior to the second dose. Restoring the receptor-TYK2 complex (RTC) partially recovered both ISGs expression, while restoring TYK2 alone had no effect (SI Figure S12), indicating that negative feedback primarily targets the RTC. Reintroducing JAK, whose levels remained unchanged across conditions, also had no impact on ISG expression. These results confirm that desensitization is driven by the depletion or inhibition of the receptor-kinase complex, which serves as the key regulator of downstream IFN signaling.

### Harnessing Interferon and Antiviral Synergies to Enhance Viral Suppression

Building on our insight about IFN-mediated immune response regulation, we examined how interferon administration can modulate the outcome of viral infection. The timing and dosage of IFN treatment are suggested as critical factors influencing its efficacy [64, 65, 66]. Our model shows that IFN treatment markedly decreases viral load in a dose-dependent fashion when given before or shortly after infection (Figure 5a). The timing of administration significantly impacts treatment efficacy. Specifically, a low IFN dose (0.3 nM) given 1–7 days before infection yields an antiviral effect over 100 times greater than a high dose (*>*10 nM) applied a few hours post-infection (Figure 5a). We observe that IFN potency declines rapidly once replication is established, with inhibition strongest during the earliest stages of infection. This is consistent with studies showing that early replication and translation events represent a vulnerable bottleneck for viral control [64], that dengue virus is sensitive to IFN only before productive replication [65], and that SARS-CoV-2 becomes largely refractory to IFN following initial replication due to antagonism of the JAK–STAT pathway [66].

**Figure 5.**
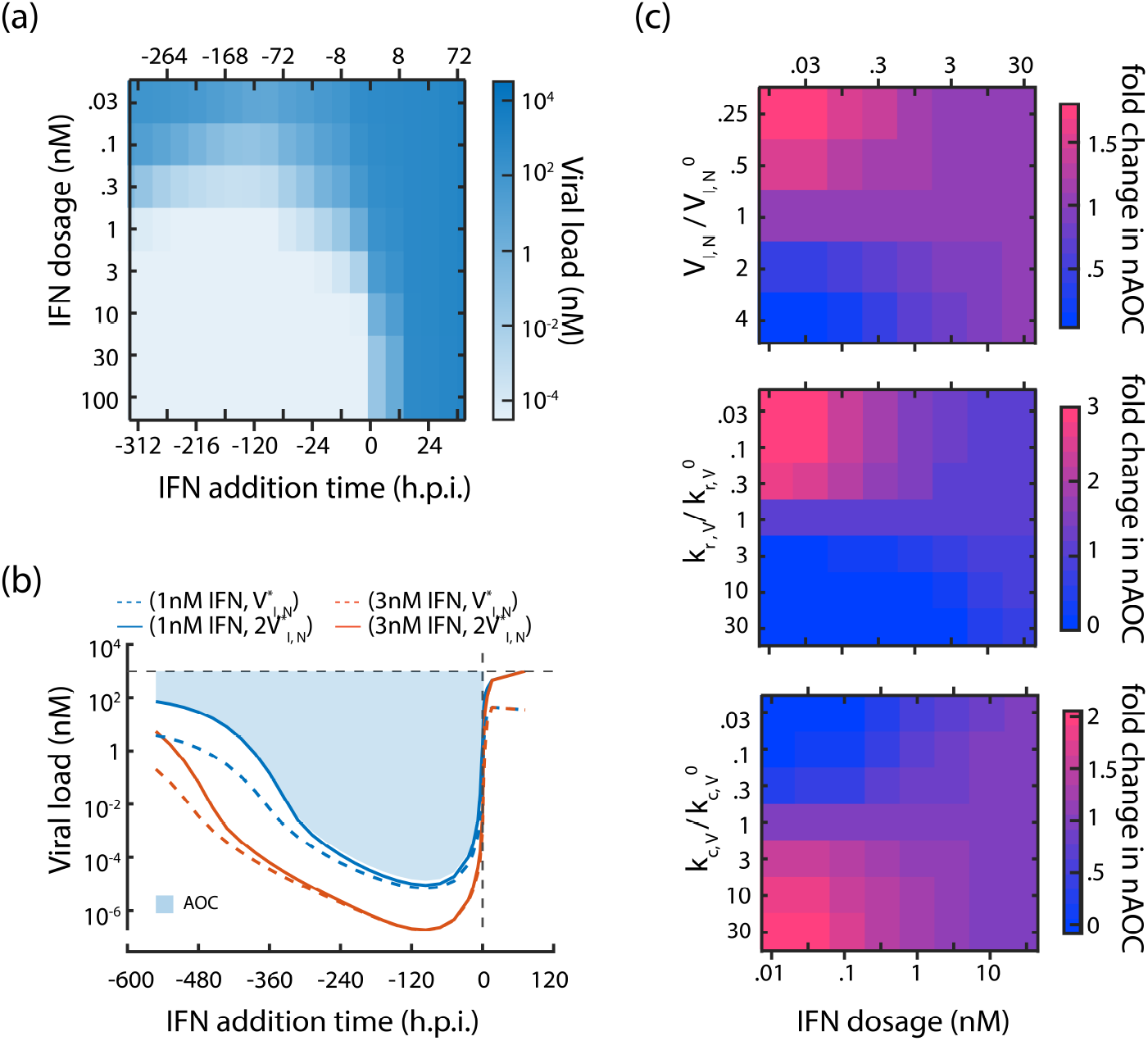
Effect of IFN dosage and timing on viral infection. (a) (a) Heatmap showing the viral load (nM) as a function of the timing of IFN addition (hours postinfection, h.p.i.) and the IFN dosage (nM). Simulations were conducted under a virus-high regime, where the virus exhibits enhanced immune escape capabilities (virus-high regime; 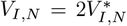). (b) Viral load dynamics over time of IFN treatment for two IFN dosages (blue: 1 nM, red: 3 nM) and two levels of viral countermeasure efficiency (dotted lines: 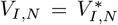; solid lines: 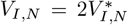) are shown. The prophylactic efficacy is quantified using the area over the curve (AOC), represented by the light blue shaded region. A higher AOC value corresponds to a stronger prophylactic effect, as it reflects a greater reduction in viral load. (c) Heatmaps illustrating the fold change in normalized AOC (nAOC) as a function of IFN dosage (nM) and key viral parameters: (top) viral countermeasure efficiency (*V*_*I,N*_ ), (middle) viral replication rate (*k*_*r,V*_ ) and (bottom) viral compartmentalization rate (*k*_*c,V*_ ). Warmer colors (pink) indicate higher fold changes in nAOC, signifying improved prophylactic efficacy.

Prompted by this result, we examine the effectiveness of IFN as a prophylactic agent and its efficacy in combination with direct-acting antivirals (Figure 5c and SI Figure S13). To fairly compare the effect of IFN exposure to viral infection, we quantify antiviral efficacy by estimating the fold reduction in viral load due to IFN exposure (or combination therapy with antivirals), integrated over the course of IFN exposure (area-over-curve metric, AOC; Figure 5b). This area-over-curve metric, normalized against no IFN exposure (referred here as nAOC, see Methods), captures the time-of-addition-independent antiviral efficacy of the strategy, with higher nAOC scores indicating greater efficacy of antiviral treatment.

AOC for a low IFN exposure of 1 nM (*≈*132, dashed blue line, Figure 5b) is less than that of a higher dose exposure of 3 nM (*≈*175, dashed red line, Figure 5b). Similarly, AOC of 1 nM prophylactic IFN exposure is higher when the viral antagonism (*V*_*I,N*_ ) is low; AOC = 132 when 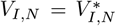 (dashed blue line, Figure 5b) compared to AOC = 115 when 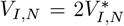 (solid blue line, Figure 5b). These observations are expected as the efficacy of IFN exposure should generally increase with IFN dosage and decrease with the strength of viral antagonism.

We next investigate how direct-acting antivirals (DAAs) can enhance the prophylactic efficacy of IFN exposure. The enhancement is estimated as the fold change in nAOC value due to the perturbation of viral processes in the presence of IFN prophylaxis. We observe an enhanced efficacy in the reduction of viral load when a prophylactic IFN treatment with direct-acting antivirals (DAAs) that limit viral antagonism (Figure 5c, top; SI Figure S13b), reduce viral replication (Figure 5c, middle), or enhance the formation of compartmentalized replication complexes (Figure 5c, bottom). Interestingly, we observe that the enhancement in antiviral efficacy is higher when either the IFN dosage is low or the strength of DAAs is higher, that is, when 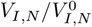 or 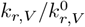 is lower or 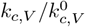 is higher. Overall, these findings underscore the importance of early IFN exposure in prophylaxis, especially in combination with direct-acting antivirals, and demonstrate how synergy between antiviral drugs and innate immunity can be leveraged to enhance antiviral action.

## Discussion

By integrating representations of the viral life cycle with innate immune signaling pathways, our approach provides a framework to analyze the coupled dynamics of virus-host interactions. Specifically, the inclusion of the RIG-I-IFN and JAK-STAT signaling pathways, along with dsRNA-induced immune activation and viral immune evasion mechanisms, allows us to examine the feedback loops and molecular interactions that drive these dynamics. Our model suggests bifurcation points that determine infection outcomes, transitioning between “virus-high, ISG-low” and “virus-low, ISG-high” states, governed by thresholds in viral antagonism and immune antiviral activity. These thresholds reveal an inverse relationship between viral load and ISG levels, as observed in single-cell studies. Similar bimodal infection have been observed in other single-cell experiments [29, 67, 41, 67, 44, 68]. For example, O’Neal et al. [67] demonstrated that in West Nile virus-infected cells, ISG expression declined with increasing viral abundance due to antagonism of the IFN-I signaling. While such “all-or-nothing” ISG induction has been attributed to cell-intrinsic noise previously [29], our model suggests that both viral and host factors shape the infection outcomes. This is consistent with studies on IFN-I dynamics [41, 44, 28], which link variability in viral replication, IFN-I signaling, and amplification capacity to the all-or-nothing induction of antiviral states.

By incorporating viral dsRNA dynamics and their role in triggering RIG-I signaling, our model demonstrates the balance between viral replication and immune detection and emphasizes the importance of early immune activation. RIG-I-mediated responses, as observed in JEV, are driven by the kinetics of dsRNA compartmentalization and the availability of replication intermediates in the cytoplasm [51, 69]. In contrast, we propose that coronaviruses shield dsRNA effectively, delaying immune detection and enabling viral propagation [70, 71, 72]. This is in agreement with research showing that interfering with replication compartments or viral mutants that have impaired shielding lead to increased dsRNA leakage, which boosts immune activation and assists in viral clearance[73, 74, 75]. Our analysis identifies two strategies used by (+)RNA viruses to balance replication and immune evasion. *V irulent Replicators*, such as poliovirus and JEV, produce dsRNA rapidly, triggering immune responses but countering this with immune suppression to complete replication before clearance. On the other hand, *Latent Replicators*, such as HCV and dengue virus, replicate more slowly but effectively shield dsRNA within membranous compartments to evade detection and establish chronic infections. We propose that these strategies reflect a trade-off between replication and immune evasion. Our model offers a framework to differentiate these viral strategies and analyze how the viral life cycle and immune evasion influence infection outcomes.

We identified three distinct phases in the antiviral immune response: an early sensing and activation phase, an intermediate signaling phase, and a late resolution phase. Early responders like aMAVS, pIKK, and pTBK1 are rapidly induced by viral RNA sensing through RIG-I, consistent with known RLR signaling mechanisms [9]. These activation markers act as proximal hubs facilitating downstream signaling, with their timing being critical for an effective response to RNA viruses. For instance, activated MAVS aggregates form antiviral signalosomes providing a scaffold for sustained signaling that sustains prolonged IRF3/7 and NF*κ*B activation, [76, 77]. Likewise, TBK1 and IKK*ϵ* phosphorylation serves as a critical early checkpoint, connecting MAVS activation to IRF3/7 phosphorylation and the induction of interferon genes [78]. In the intermediate phase, IRF and IFN-related factors drive the expression of antiviral ISGs, although this response is suppressed under high viral loads. During the late phase, IFN signaling through JAK-STAT components becomes prominent but is often attenuated in severe infections due to feedback inhibition or impaired upstream signaling [50, 69]. These temporally distinct phases help identify key regulators at each stage, providing a framework to prioritize therapeutic targets based on the timing and nature of immune engagement. For instance, we observed that phosphorylated IRF7, nascent IFN transcripts, and early ISGs were consistently different between mild and severe infections early in the response, making them strong predictors of infection outcomes. This is supported by experimental evidence showing that IRF7 is rapidly phosphorylated upon viral sensing, driving the second wave of type I IFN production essential for antiviral defense. IRF7 deficiency severely impairs early IFN responses, underscoring its critical role in immune activation [79, 80, 81]. Consistent with our model’s predictions, type I interferons, particularly interferon *β* (IFN-*β*), are recognized as markers of effective immune response, where reduced peripheral IFN-*β* levels during SARS-CoV-2 infection is associated with the severity of the disease [82]. Thus, these temporally resolved immune effector modules elucidate the complexity of immune models and identify mechanistic checkpoints at different phases.

A significant outcome from our molecular-level analysis is the capacity to replicate IFN desensitization upon repeated IFN exposure in a dose-dependent manner. Our model identifies IFN receptor–associated kinases, particularly TYK2 and Receptor-TYK2 Complex (RTC), as key nodes responsible for IFN-desensitization. Previously, it was shown that TYK2 is essential for type 1 IFN signaling, leading to STAT pathway-mediated activation of negative regulators such as the Suppressor of Cytokine Signaling (SOCS) and USP18 [9, 58, 62, 63, 83, 84, 85]. Prolonged or repeated exposure to interferons diminishes receptor availability and promotes SOCS-mediated enhances SOCS-driven degradation of the receptor-tyrosine complex (RTC). Consistent with this, we propose that the receptor replenishment dynamics via RTC are pivotal in limiting IFN signaling, leading to IFN desensitization after substantial exposure.

We also show that low-dose prophylactic administration of IFN can considerably inhibit HCV replication, surpassing the effectiveness of treatments post-infection. This is consistent with previous studies on (+)RNA viruses like SARS-CoV-2 and dengue, where delayed IFN treatment was shown to be ineffective or even detrimental in humans and cell culture experiments [64, 66, 65]. Our findings contribute to a growing body of work suggesting that the timing and dosage of immune interventions can strongly influence disease outcomes [86, 87]. Further, we demonstrate how model-informed combination therapies, including those targeting viral processes and inherent antagonistic feedback loops, can significantly enhance IFN prophylaxis. In fact, pairing IFN-*α*2a with direct-acting antivirals like nucleoside analogues (for example, remdesivir or EIDD-2801) or agents targeting the host (such as camostat or cycloheximide) has demonstrated considerable enhancement of IFN-induced antiviral effects and improved viral inhibition in both in vitro and in vivo studies [88]. Similar evidence from human trials with intranasal prophylactic administration of IFN-*α* significantly reduced the incidence and severity of rhinovirus and coronavirus infections, while inconsistent benefits were reported for post-exposure prophylaxis [89]. Thus, interventions employing IFN prophylaxis, especially in high-risk populations or during outbreaks, might be a viable strategy.

To keep our model tractable while capturing the essential details of the host-virus interactions, we have made tactical assumptions and simplifications. Our model lumps the sensing by pattern recognition receptors (PRRs) and assumes an averaged cellular activation. Although multiple classes of PRR (TLRs, RLRs, NLRs, CLRs) are involved in shaping innate immune responses, it is well established that viral dsRNA is a major trigger for antiviral responses [90, 91]. Consequently, we limit ourselves to dsRNA-sensing pathways to emulate RNA virus immune detection. To simplify the interpretation of innate immune feedback, we also demonstrate system behavior with a uniform level of negative feedback across all signaling nodes and ignore the strength of context-dependent regulators (e.g., SOCS activity on RTC), which can vary by cell type and stimulus. However, our model enables explicit control of molecular-level interactions through independently defined parameters. We observe that individual molecular processes also exhibit bifurcation between ‘virus-high, ISG-low’ and ‘virus-low, ISG-high’ states suggesting this to be a system-wide feature. In the future, the model can be used to explicitly investigate the role of these interactions. We modeled antivirals as reductions in viral parameters, though incorporating drug pharmacodynamics could improve outcome predictions. Despite this simplification, our framework provides a robust basis for studying molecular-specific antiviral responses with flexibility for refinement. Since our model can capture molecular interactions, we can incorporate cellular and tissue heterogeneity. Combined with spatio-temporal dynamics and multi-cell immune responses, our approach can be extended to study infection dynamics in complex tissues and organs.

We present a universal framework for understanding virus-host dynamics and identifying therapeutic targets. The non-linear dependence of viral load on parameters like viral antagonism and immune activation reveals critical points for shifting outcomes toward viral clearance. This framework supports the design of model-informed antiviral strategies while highlighting the importance of early immune activation, ISG dynamics, and IFN signaling in balancing viral persistence and immune control.

## Methods

### Models integration and extension

To create a detailed, mechanistic model of the virus–innate immune response, we integrated the detailed intracellular viral life cycle model by Chhajer *et al*. [24] with the RIG-I–IFN–ISG signaling model by Burkart *et al*. [39]. To achieve this integration, we introduced additional variables, parameters, and equations to capture extracellular viral dynamics, the leakage and sensing of double-stranded RNA (dsRNA), the regulation of Interferon Regulatory Factor 7 (IRF7), and multilayered feedback mechanisms involving both viral antagonism and host regulatory pathways.

The viral life cycle model was extended to include extracellular dynamics of viral entry and internalization. Viral entry was modeled as a first-order process governed by the rate constant *k*_en_, leading to depletion of extracellular virus (*V*_0_). Internalized virus (*V*_*I*_ ) accumulates as a result and can either undergo uncoating at a rate *k*_*f,V*_ or be degraded at a rate 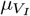. Upon uncoating, viral RNA is released into the cytoplasm and contributes to the cytoplasmic viral RNA pool (*R*_cyt_). To account for this, the original equation describing cytoplasmic viral RNA dynamics was modified to include an additional term reflecting input from uncoated virus. All other terms in this equation, as well as the ODEs describing the dynamics of structural and non-structural viral proteins (*P*_*S*_, *P*_*NS*_), replication compartment formation (*RC*_*form_rate*_), and replication complexes within compartments (*RC*_*CM*_ ), remain identical to the original framework by Chhajer *et al*. [24].

A critical extension of the viral model was the explicit inclusion of dsRNA leakage from replication compartments. While the original model did not accommodate this, cytosolic dsRNA serves as a key activator of the innate immune response and is central to coupling viral replication with RIG-I sensing. We therefore introduced a leakage term at rate *k*_*l,V*_, following an approach similar to Zitzmann *et al*. [32]. *Once in the cytoplasm, dsRNA undergoes natural degradation at a rate µ*_*r,V*_, can be replenished through the deactivation of RIG-I at a rate *b*_*RIGI*_, and is cleared through RIG-I–mediated sensing at a rate proportional to the product of RIG-I and dsRNA concentrations (*k*_RIGI_ *·* RIGI *·* RNA_*ds*_). This formulation establishes the mechanistic link between the viral life cycle and the innate immune module.

The innate immune signaling model of Burkart *et al*. [39] was extended by incorporating the dynamics of Interferon Regulatory Factor 7 (IRF7). IRF7 is a transcription factor essential for the induction of type I interferon genes, particularly in plasmacytoid dendritic cells and during late-phase antiviral responses [92]. It is also responsible for its own induction via IFN-dependent transcription [93]. Given its high degree of homology with IRF3, we assumed IRF7 undergoes phosphorylation and nuclear translocation through a mechanism analogous to IRF3. Specifically, cytoplasmic IRF7 is phosphorylated by activated TBK1 and IKK*ϵ*, translocates into the nucleus, and regulates interferon gene expression. Once in the nucleus, phosphorylated IRF7 (pIRF7) can be dephosphorylated or exported back into the cytoplasm. Its synthesis is driven by translation from IFN-induced mRNA, while constitutive degradation occurs with a first-order decay rate.

In addition to coupling viral and immune processes mechanistically, we introduced multilayered feedback to capture the dynamic regulation that occurs at different levels. On the host side, interferon-stimulated genes (ISGs) target distinct stages of the viral life cycle. For example, CH25H and IFITMs inhibit viral entry [9, 12], IFIT and TRIM family proteins disrupt RNA and protein synthesis [9, 12],

and viperin interferes with assembly and egress by disrupting viral envelope formation [9, 12]. In contrast, viruses of the *Flaviviridae* family have evolved to counteract host defenses by antagonizing RIG-I signaling and JAK-STAT signaling cascades, thereby suppressing type I IFN production and signaling [48, 49, 50]. Finally, the host itself incorporates intrinsic negative regulatory mechanisms to limit excessive immune activation. In particular, molecules such as SOCS proteins and USP18 dampen IFN-induced signaling, ensuring resolution of the antiviral state [61, 62, 13].

To quantitatively capture these interacting layers of feedback, we introduced three lumped pseudo- variables representing distinct functional modules: *V*_*I,N*_ (efficacy of viral antagonism), *I*_*V,N*_ (potency of ISG-mediated antiviral activity), and *I*_*I,N*_ (efficiency of host negative regulation of immune signaling). These pseudo-variables act by modulating either the intrinsic process rates or the degradation rates of relevant molecular species. A complete list of all model equations, including the ones mentioned below (SI-Model equations), along with detailed variable definitions, initial conditions (tables S1-S2), parameter values, and sources (tables S3-S7), is provided in the Supplementary Information.

#### Viral life cycle and dsRNA detection

We added the following variables and equations to the viral life cycle model proposed by Chhajer et al.[24] to incorporate the extracellular viral dynamics of viral entry and internalization:

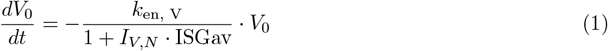

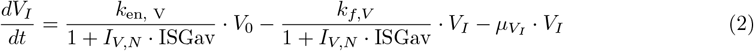

Subsequently, the equation monitoring the levels of cytoplasmic viral RNA is changed:

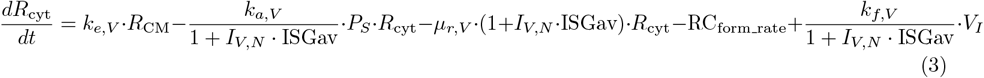

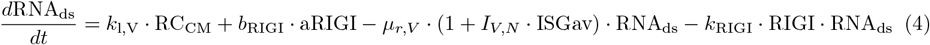

#### Innate immune signaling pathway

The dynamics of Interferon Regulatory Factor 7 (IRF7) are modelled as shown below:

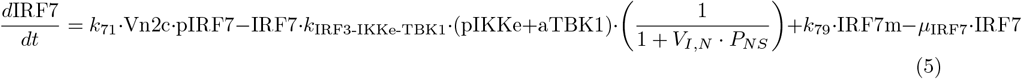

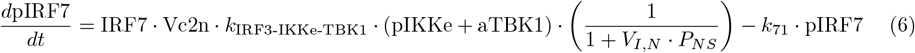

#### Incorporation of feedback at multiple levels by both the virus and innate immune system

Feedback was modeled as a generic modulation of rates:

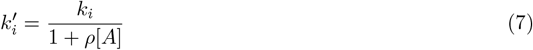

and the degradation rates *µ*_*i*_ are changed as

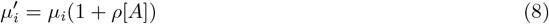

where *k*_*i*_ denotes rate constants of viral entry (*k*_*en,V*_ ), fusion (*k*_*f,V*_ ), replication (*k*_*r,V*_ ), translation (*k*_*t,V*_ ), egress (*k*_*a*_), and *µ*_*i*_ represents degradation rates of viral RNA (*µ*_*r,V*_ ), and proteins (*µ*_*p,V*_ ), *ρ* is *I*_*V,N*_ and [*A*] is levels of antiviral ISG (ISGav). For immune signaling processes, *ρ* = *V*_*I,N*_, with viral proteins (*P*_*NS*_) attenuating MAVS activation (*k*_MAVS_), IRF3/IRF7 activation (*k*_*IRF* 3_*IKKe_T BK*1_), and STAT complex activation (*k*_13_). In addition, host-driven negative feedback is incorporated by setting *ρ* = *I*_*I,N*_ and [*A*] = ISGn, which downregulates activated receptor complex (ARC) formation (*k*_7_).

### Calculation of critical values and bifurcation point sensitivity analysis

To identify the critical value of 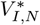(or 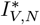), we systematically vary one of *V*_*I,N*_ or *I*_*V,N*_ keeping the other fixed, and record the extracellular viral concentration at 48 hours post-infection (*V*_*T*_ ). Next, we evaluate the first-order numerical derivative of the logarithm of viral output (log_10_(*V*_*T*_ )) with respect to the variable varied and identify the point of maximum change as the critical value. *V*_*I,N*_ and *I*_*V,N*_ were varied in the logarithm scale to span a large dynamic range.

To evaluate how different model parameters influence the critical values of immune activity and viral antagonism, we performed Bifurcation Sensitivity Analysis. To evaluate the sensitivity of 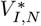 at the bifurcation point, ‘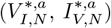’, we fix *I*_*V,N*_ as 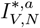. When we look at the ‘*i*^*th*^’ model parameter, we alter the value of the parameter from its nominal value 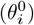 to *θ*_*i*_ and re-evaluate the critical value of viral antagonism 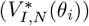. Finally, we evaluate the sensitivity index as the ratio of the relative change in threshold value with respect to the relative change in the parameter value, using the following equation:

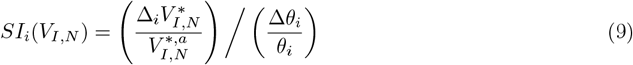

As sensitivity captures the slope of the threshold function with respect to the model parameter, we use a five-point central difference formula instead of the naive single-point difference formula, to calculate the sensitivity index more accurately. The threshold values were evaluated at 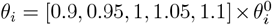.

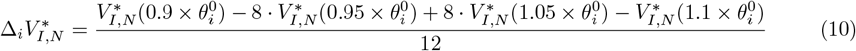

*SI*_*i*_(*V*_*I,N*_ ) determines the influence of the ‘*i*^*th*^’ model parameter on the critical viral antagonism required for immune escape. Analogously, we can calculate *SI*_*i*_(*I*_*V,N*_ ); here we fix *V*_*I,N*_ as 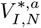, and assess how changes in the model parameters affect the immune activity threshold 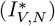 necessary for viral suppression. To identify the most important parameters that influence both the thresholds, we rank the parameters based on threshold sensitivity score, calculated as the sum of squares of the corresponding sensitivity indices, i.e., (*SI*_*i*_(*V*_*I,N*_ ))^2^ + (*SI*_*i*_(*I*_*V,N*_ ))^2^

### Latin Hypercube Sampling and correlation with viral load

To evaluate the influence of host and viral parameters on infection outcomes, we performed Latin Hypercube Sampling (LHS). Each model parameter *p*_*i*_ was varied within a log-uniform range

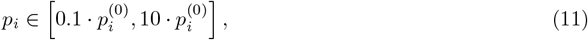

where 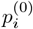 denotes the original value of parameter *i*. A total of *N* = 25000 parameter sets were generated, each comprising the full parameter vector,

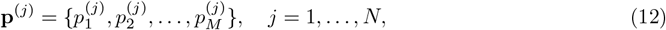

To quantify the relationship between individual parameters and infection outcome, we computed the Pearson correlation coefficient between each parameter *p*_*i*_ and the viral load across all LHS-generated parameter sets:

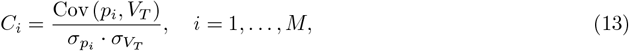

where *σ*_*p*_*i* and *σ*_*V,T*_ denote the standard deviations of parameter *p*_*i*_ and viral load *V*_*T*_, respectively. The resulting correlation coefficients *C*_*i*_ were then ranked to identify the top 10 positively correlated and top 10 negatively correlated parameters. These parameters were visualized in a bar plot to highlight the most influential drivers of viral load variability (Figure 2d).

### Normalized dynamics based clustering

In our analysis, we perform hierarchical clustering on normalized time-series data for each model variable under three conditions - A: low *V*_*T*_ (when 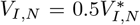), B: critical *V*_*T*_ (when 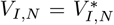), and C: high *V*_*T*_ (when 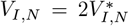 ). Variables with zero standard deviation or with a low coefficient of variation 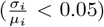 were excluded, ensuring retention of only dynamically informative variables. We then concatenate the dynamics of the selected variables in the three regimes and normalize them using Z-score normalization. The mean and standard deviation used for the normalization is based on the concatenated dynamics.

A pairwise distance matrix was computed across all retained variables, followed by hierarchical clustering based on the average linkage criterion. The resulting dendrogram was cut to yield k=8 clusters. For each cluster, variable memberships and their average correlation to the cluster mean trajectory were computed. Variable-wise dynamics were additionally annotated with marker sizes inversely proportional to the distance to the centroid.

This analysis was based on our recently published model-based experiment design algorithm, PARSEC [94]. Like the analysis here, the PARSEC groups model variables/observables, but the clustering is based on differences in parameter sensitivity signatures instead of dynamical correlations.

### Quantification of dynamic differences using a *χ*^2^-like trajectory score

To quantify dynamic differences in temporal responses between conditions (specify the conditions), we defined a log-transformed mean squared deviation (*χ*^2^-like) score for each variable. For a given variable *x*_*k*_, the score between two conditions *c*_1_ and *c*_2_ was calculated as:

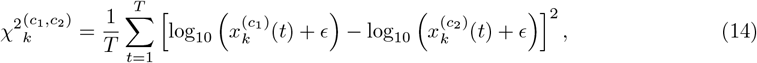

where 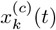 denotes the trajectory of variable *k* at time *t* under condition *c, T* is the number of time points. This measure captures the average squared deviation between log-scaled trajectories and is mathematically equivalent to a *χ*^2^-like score. A small positive number *ϵ* = 10^*−*8^ is added to the variables so that the resulting logarithmic value is defined and finite.

We applied this metric in two distinct contexts. First, in the clustering analysis, it was used to compare responses between conditions A and C (Figure 3c). Second, in the interferon (IFN) stimulation experiments, the same metric was employed to compare trajectories between two IFN priming protocols: one in which cells received 100 nM IFN at 24 h following a 1 nM priming dose, and another in which the same 100 nM boost followed a 100 nM priming dose (Figure 4c).

### Normalized area over the curve (nAOC) calculation - estimating prophylactic efficiency of IFN pretreatment

To evaluate the prophylactic action, we simulate the effect of various dosages of IFN (as shown in Figure administered *k* days before the start of infection, where *k ∈* [0, 1, 2, … 30]. We report the resulting viral load at 48 hours post-infection (*V*_*T*_ (*k*)). In the logarithmic scale, we compute the area between this curve (log10(*V*_*T*_ (*t*))) and a horizontal line at *y*_*th*_ = 3 (a high viral load threshold reference) between *k* = 0 to *k* = 30 days using the trapezoidal rule. Specifically, the AOC was calculated using:

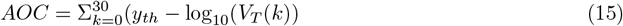

## Availability of code

Codes for simulation and analysis can be downloaded from our GitHub repository, https://github.com/ramyab-986/Integrated_Virus-Immune_Model/tree/master.

## Ethics Statement

This is a computational study. No animals or humans and/or their tissue samples were used.

## Acknowledgement

We thank Mahammed Kaif, Muhsin Chalil, and Suraj Jagtap for valuable feedback on the manuscript.

## Author Contributions

Data extraction: RB. Model development, software, and formal analysis: RB, HC. Conceptualization, result interpretation, and original draft preparation: RB, HC, RR.

## Funding

This work was supported by the Indian Institute of Science, Bangalore (RR), Wellcome Trust—DBT India Alliance intermediate fellowship (RR).

## Competing interests

The authors declare that they have no competing interests.

## Notes

### Competing Interest Statement

The authors have declared no competing interest.

### Summary of Updates

The text and figures were revised.

