## Supplementary material for "Integrative Modelling of Innate Immune Response Dynamics during Virus Infection": SI

Orchid identifiers: RB: 0000-0002-0056-9980; HC: 0000-0001-9252-1855; RR: 0000-0003-3329-8803

[c] Equal contribution.

*Corresponding author: Rahul Roy.

Department of Chemical Engineering, Indian Institute of Science, Bangalore, Karnataka, India-560012.

Article classification: Biological Sciences | Microbiology and Immunology

Keywords: Positive-sense RNA virus | Innate immune signaling | Mathematical modeling | Interferon therapy | Virus-host dynamics

### Supplementary Information

Integrative Modelling of Innate Immune Response Dynamics during Virus Infection

Ramya Boddepalli, Harsh Chhajer and Rahul Roy

#### Model Equations

$$f_{CM} = 1 - \exp\left(-\left(\frac{t}{\tau}\right)^4\right)$$

$$RC_{\text{form\_rate}} = k_{c,V} \cdot R_{\text{cyt}} \cdot P_{NS} \cdot \left(f_{CM} - \frac{RC_{CM}}{N_C}\right)$$

$$\frac{dV_0}{dt} = -\frac{k_{\text{en, V}}}{1 + I_{V,N} \cdot \text{ISGav}} \cdot V_0 \qquad \text{(eq S1)}$$

$$\frac{dV_T}{dt} = \frac{k_{a,V}}{1 + I_{V,N} \cdot \text{ISGav}} \cdot P_S \cdot R_{\text{cyt}} - \mu_{V,V} \cdot V_T \qquad \text{(eq S2)}$$

$$\frac{dV_I}{dt} = \frac{k_{\text{en, V}}}{1 + I_{V,N} \cdot \text{ISGav}} \cdot V_0 - \frac{k_{f,V}}{1 + I_{V,N} \cdot \text{ISGav}} \cdot V_I - \mu_{V_I} \cdot V_I \qquad \text{(eq S3)}$$

$$\frac{dR_{\text{cyt}}}{dt} = k_{e,V} \cdot R_{\text{CM}} - \frac{k_{a,V}}{1 + I_{V,N} \cdot \text{ISGav}} \cdot P_S \cdot R_{\text{cyt}} - \mu_{r,V} \cdot (1 + I_{V,N} \cdot \text{ISGav}) \cdot R_{\text{cyt}} - \text{RC}_{\text{form\_rate}} + \frac{k_{f,V}}{1 + I_{V,N} \cdot \text{ISGav}} \cdot V_I \qquad \text{(eq S4)}$$

$$\frac{dR_{\text{CM}}}{dt} = \frac{k_{r,V}}{1 + I_{V,N} \cdot \text{ISGav}} \cdot \text{RC}_{\text{CM}} - k_{e,V} \cdot R_{\text{CM}} \qquad \text{(eq S5)}$$

$$\frac{dP_S}{dt} = \frac{k_{t,V}}{1 + I_{V,N} \cdot \text{ISGav}} \cdot R_{\text{cyt}} - \frac{k_{a,V}}{1 + I_{V,N} \cdot \text{ISGav}} \cdot \text{nSP} \cdot P_S \cdot R_{\text{cyt}} - \mu_{p,V} \cdot (1 + I_{V,N} \cdot \text{ISGav}) \cdot P_S \qquad \text{(eq S6)}$$

$$\frac{dP_{\text{NS}}}{dt} = \frac{k_{t,V}}{1 + I_{V,N} \cdot \text{ISGav}} \cdot R_{\text{cyt}} - \text{RC}_{\text{form\_rate}} - \mu_{p,V} \cdot (1 + I_{V,N} \cdot \text{ISGav}) \cdot P_{\text{NS}} \qquad \text{(eq S7)}$$

$$\frac{d\text{RC}_{\text{CM}}}{dt} = \text{RC}_{\text{form\_rate}} - k_{\text{l,V}} \cdot \text{RC}_{\text{CM}} \qquad \text{(eq S8)}$$

$$\frac{d\text{RNA}_{\text{ds}}}{dt} = k_{\text{l,V}} \cdot \text{RC}_{\text{CM}} + b_{\text{RIGI}} \cdot \text{aRIGI} - \mu_{r,V} \cdot (1 + I_{V,N} \cdot \text{ISGav}) \cdot \text{RNA}_{\text{ds}} - k_{\text{RIGI}} \cdot \text{RIGI} \cdot \text{RNA}_{\text{ds}} \qquad \text{(eq S9)}$$

$$\frac{d\text{RIGI}}{dt} = k_{\text{RIGI\_synt}} - \mu_{\text{RIGI}} \cdot \text{RIGI} + b_{\text{RIGI}} \cdot \text{aRIGI} - k_{\text{RIGI}} \cdot \text{RIGI} \cdot \text{RNA}_{\text{ds}} + k_{\text{t, ISG RNA}} \cdot \text{ISGav}_{\text{m}} \qquad \text{(eq S10)}$$

$$\frac{d\text{aRIGI}}{dt} = k_{\text{RIGI}} \cdot \text{RIGI} \cdot \text{RNA}_{\text{ds}} - \mu_{\text{RIGI}} \cdot \text{aRIGI} - b_{\text{RIGI}} \cdot \text{aRIGI} \tag{eq S11}$$

$$\frac{d\text{MAVS}}{dt} = b_{\text{MAVS}} \cdot \text{aMAVS} - \text{MAVS} \cdot \text{aRIGI} \cdot \left(\frac{k_{\text{MAVS}}}{1 + V_{I,N} \cdot P_{\text{NS}}}\right) \tag{eq S12}$$

$$\frac{d\text{aMAVS}}{dt} = \text{MAVS} \cdot \text{aRIGI} \cdot \left(\frac{k_{\text{MAVS}}}{1 + V_{I,N} \cdot P_{\text{NS}}}\right) - b_{\text{MAVS}} \cdot \text{aMAVS} \tag{eq S13}$$

$$\frac{d\text{IKKe}}{dt} = b_{\text{KINASE}} \cdot \text{aIKKe} - \text{IKKe} \cdot \text{aMAVS} \cdot k_{\text{IKKe-TBK1}} \tag{eq S14}$$

$$\frac{d\text{aIKKe}}{dt} = \text{IKKe} \cdot \text{aMAVS} \cdot k_{\text{IKKe-TBK1}} - b_{\text{KINASE}} \cdot \text{aIKKe} \tag{eq S15}$$

$$\frac{d\text{TBK1}}{dt} = b_{\text{KINASE}} \cdot \text{aTBK1} - \text{TBK1} \cdot \text{aMAVS} \cdot k_{\text{IKKe-TBK1}} \tag{eq S16}$$

$$\frac{d\text{aTBK1}}{dt} = \text{TBK1} \cdot \text{aMAVS} \cdot k_{\text{IKKe-TBK1}} - b_{\text{KINASE}} \cdot \text{aTBK1} \tag{eq S17}$$

$$\frac{d\text{IRF3}}{dt} = \text{Vn2c} \cdot b_{\text{IRF3}} \cdot \text{pIRF3} - \text{IRF3} \cdot k_{\text{IRF3-IKKe-TBK1}} \cdot (\text{pIKKe} + \text{aTBK1}) \cdot \left(\frac{1}{1 + V_{I,N} \cdot P_{\text{NS}}}\right) \tag{eq S18}$$

$$\frac{d\text{pIRF3}}{dt} = (\text{IRF3} \cdot \text{Vc2n} \cdot k_{\text{IRF3-IKKe-TBK1}} \cdot (\text{pIKKe} + \text{aTBK1})) \cdot \left(\frac{1}{1 + V_{I,N} \cdot P_{\text{NS}}}\right) - b_{\text{IRF3}} \cdot \text{pIRF3} \tag{eq S19}$$

$$\frac{d\text{IKK}}{dt} = \text{aIKK} \cdot b_{\text{KINASE}} - \text{IKK} \cdot \text{aMAVS} \cdot k_{\text{IKK}} \tag{eq S20}$$

$$\frac{d\text{aIKK}}{dt} = \text{IKK} \cdot \text{aMAVS} \cdot k_{\text{IKK}} - \text{aIKK} \cdot b_{\text{KINASE}} \tag{eq S21}$$

$$\frac{d\text{NFkB\_IkBa}_c}{dt} = \text{IkBac} \cdot \text{NFkBc} \cdot k_{\text{inh\_p65}} - \text{NFkB\_IkBac} \cdot \text{aIKK} \cdot k_{\text{act}} \tag{eq S22}$$

$$\frac{dp\text{NFkB}_n}{dt} = \text{Vc2n} \cdot \text{aIKK} \cdot k_{\text{act}} \cdot (\text{NFkBc} + \text{NFkB\_IkBac}) - k_{\text{deph}} \cdot p\text{NFkBn} \tag{eq S23}$$

$$\frac{d\text{NFkB}_n}{dt} = k_{\text{deph}} \cdot p\text{NFkBn} - \text{NFkBn} \cdot k_{\text{transp\_NFkB}} \quad \text{(eq S24)}$$

$$\frac{d\text{NFkB}_c}{dt} = \text{NFkBn} \cdot \text{Vn2c} \cdot k_{\text{transp\_NFkB}} - \text{IkBac} \cdot \text{NFkBc} \cdot k_{\text{inh\_p65}} - \text{NFkBc} \cdot \text{aIKK} \cdot k_{\text{act}} \quad \text{(eq S25)}$$

$$\frac{d\text{IkBa}_c}{dt} = k_{\text{expr\_IkBa}} \cdot (\text{NFkBn} + p\text{NFkBn}) - \text{IkBac} \cdot \mu_{\text{IkBa}} - \text{IkBac} \cdot \text{NFkBc} \cdot k_{\text{inh\_p65}} \quad \text{(eq S26)}$$

$$\frac{d\text{IFNb\_}m}{dt} = \text{B}_O \cdot \text{Vn2c} \cdot k_{\text{TFBS\_IFNb}} - \text{IFNb}_m \cdot \mu_{\text{m\_IFNb}} + \text{Vn2c} \cdot k_{\text{m\_IFNb}} \cdot (p\text{IRF3} + p\text{IRF7}) \cdot (\text{NFkBn} + p\text{NFkBn}) \quad \text{(eq S27)}$$

$$\frac{d\text{IFNa\_}m}{dt} = \text{B}_O \cdot \text{Vn2c} \cdot k_{\text{TFBS\_IFNa}} - \text{IFNa}_m \cdot \mu_{\text{m\_IFNa}} + \text{Vn2c} \cdot k_{\text{m\_IFNa}} \cdot (p\text{IRF3} + p\text{IRF7}) \quad \text{(eq S28)}$$

$$\frac{d\text{IFNl\_}m}{dt} = \text{B}_O \cdot \text{Vn2c} \cdot k_{\text{TFBS\_IFNl}} - \text{IFNl}_m \cdot \mu_{\text{m\_IFNl}} + \text{Vn2c} \cdot k_{\text{m\_IFNl}} \cdot p\text{IRF3} \quad \text{(eq S29)}$$

$$\frac{d\text{IFN}_c}{dt} = (\text{IFNa}_m + \text{IFNb}_m) \cdot k_{\text{IFN}} - k_s \cdot \text{IFN}_c - \mu_{\text{IFN}} \cdot \text{IFN}_c \quad \text{(eq S30)}$$

$$\frac{d\text{IFNl}_c}{dt} = \text{IFNl}_m \cdot k_{\text{trans\_IFNl}} - \text{IFNl}_c \cdot \mu_{\text{IFNl}} \quad \text{(eq S31)}$$

$$\frac{d\text{JAK}}{dt} = \text{ARC} \cdot k_{32} + \text{RJC} \cdot k_4 + \text{ARC} \cdot I_{I,N} \cdot \text{ISGn} \cdot k_{69} - \text{JAK} \cdot \text{IFNAR2} \cdot k_3 \quad \text{(eq S32)}$$

$$\frac{d\text{RJC}}{dt} = \text{IFNARd} \cdot k_6 - \text{RJC} \cdot k_4 + \text{JAK} \cdot \text{IFNAR2} \cdot k_3 - \text{IFNex} \cdot \text{RJC} \cdot \text{RTC} \cdot k_5 \quad \text{(eq S33)}$$

$$\begin{aligned}\frac{d\text{STAT1}_c}{dt} = {} & \text{ARC-STAT12}_c \cdot k_{12} + \text{ISGF3-CP} \cdot k_{40} - \text{STAT1}_c \cdot k_{56} + \text{PSC-CP} \cdot k_{43} \\ & - \text{ARC-STAT2}_c \cdot \text{STAT1}_c \cdot k_{11} + \text{STAT1}_n \cdot \text{Vn2c} \cdot k_{57}\end{aligned} \quad \text{(eq S34)}$$

$$\frac{d\text{CP}}{dt} = \text{ISGF3-CP} \cdot k_{39} + \text{ISGF3-CP} \cdot k_{40} + \text{PSC-CP} \cdot k_{42} + \text{PSC-CP} \cdot k_{43} - \text{CP} \cdot \text{ISGF3}_c \cdot k_{38} - \text{CP} \cdot \text{PSC}_c \cdot k_{41} \quad \text{(eq S35)}$$

$$\frac{d\text{ISGn}}{dt} = k_{transISGn} \cdot \text{ISGn\_m}_c - \text{ISGn} \cdot k_{31} \quad \text{(eq S36)}$$

$$\frac{d\text{IFNex}}{dt} = \text{IFNARd} \cdot k_6 + k_s \cdot \text{IFN}_c - \text{IFNex} \cdot \text{RJC} \cdot \text{RTC} \cdot k_5 \quad \text{(eq S37)}$$

$$\frac{d\text{STAT2}_c}{dt} = \text{ARC-STAT2}_c \cdot k_{10} + \text{ISGF3-CP} \cdot k_{40} - \text{STAT2}_c \cdot k_{58} + \text{STAT2-IRF9}_c \cdot (k_{36} + k_{61}) + \text{PSC-CP} \cdot k_{43} - \text{ARC} \cdot \text{STAT2}_c \cdot k_9 - \text{IRF9}_c \cdot \text{STAT2}_c \cdot k_{60} + \text{STAT2}_n \cdot \text{Vn2c} \cdot k_{59} \quad \text{(eq S38)}$$

$$\frac{d\text{TYK}}{dt} = \text{ARC} \cdot k_{32} + \text{RTC} \cdot k_2 + \text{ARC} \cdot I_{I,N} \cdot \text{ISGn} \cdot k_{69} - \text{IFNAR1} \cdot \text{TYK} \cdot k_1 \quad \text{(eq S39)}$$

$$\frac{d\text{ARC}}{dt} = \text{ARC\_STAT2c} \cdot k_{10} - \text{ARC} \cdot k_{34} - \text{ARC} \cdot k_{32} + \text{ARC\_STAT12} \cdot k_{13} \cdot \left(\frac{1}{1 + V_{I,N} \cdot P_{NS}}\right) + \text{IFNARd} \cdot \left(\frac{k_7}{1 + I_{I,N} \cdot \text{kinhISGn} \cdot \text{ISGn}}\right) - \text{ARC} \cdot I_{I,N} \cdot \text{ISGn} \cdot k_{69} - \text{ARC} \cdot \text{STAT2c} \cdot k_9 - \text{ARC} \cdot \text{STAT2c\_IRF9} \cdot k_8 - I_{I,N} \cdot \text{degARCISGn} \cdot \text{ISGn} \cdot \text{ARC} \quad \text{(eq S40)}$$

$$\frac{d\text{RTC}}{dt} = \text{IFNARd} \cdot k_6 - \text{RTC} \cdot k_2 + \text{IFNAR1} \cdot \text{TYK} \cdot k_1 - \text{IFNex} \cdot \text{RJC} \cdot \text{RTC} \cdot k_5 \quad \text{(eq S41)}$$

$$\frac{d\text{IFNAR1}}{dt} = \text{ARC} \cdot k_{32} + \text{RTC} \cdot k_2 + \text{ARC} \cdot I_{I,N} \cdot \text{ISGn} \cdot k_{69} - \text{IFNAR1} \cdot \text{TYK} \cdot k_1 - I_{I,N} \cdot \text{degRecISGn} \cdot \text{IFNAR1} \cdot \text{ISGn} \quad \text{(eq S42)}$$

$$\frac{d\text{IFNAR2}}{dt} = \text{ARC} \cdot k_{32} + \text{RJC} \cdot k_4 + \text{ARC} \cdot I_{I,N} \cdot \text{ISGn} \cdot k_{69} - \text{JAK} \cdot \text{IFNAR2} \cdot k_3 - I_{I,N} \cdot \text{degRecISGn} \cdot \text{IFNAR2} \cdot \text{ISGn} \quad \text{(eq S43)}$$

$$\frac{d\text{IFNARd}}{dt} = \text{ARC}\cdot k_{34} - \text{IFNARd}\cdot(k_6 + \left(\frac{k_7}{1 + I_{I,N}\cdot \text{kinhISGn}\cdot \text{ISGn}}\right)) + \text{IFNex}\cdot\text{RJC}\cdot\text{RTC}\cdot k_5 \quad \text{(eq S44)}$$

$$\frac{d\text{IRF9}_c}{dt} = k_{27} - \text{IRF9}_c\cdot(k_{29} + k_{66}) + \text{ISGF3}_c\cdot k_{15} + \text{ISGF3-CP}\cdot k_{40} + \text{STAT2-IRF9}_c\cdot k_{61} + k_{70}\cdot \text{IRF9\_m}_c$$
$$+\text{ARC}\cdot\text{STAT2-IRF9}_c\cdot k_8 - \text{IRF9}_c\cdot\text{STAT2}_c\cdot k_{60} - \text{IRF9}_c\cdot\text{PSC}_c\cdot k_{14} + \text{IRF9}_n\cdot\text{Vn2c}\cdot k_{67} \quad \text{(eq S45)}$$

$$\frac{d\text{ARC-STAT2}_c}{dt} = \text{ARC-STAT12}_c\cdot k_{12} - \text{ARC-STAT2}_c\cdot k_{10} + \text{ARC}\cdot\text{STAT2}_c\cdot k_9$$
$$+\text{ARC}\cdot\text{STAT2-IRF9}_c\cdot k_8 - \text{ARC-STAT2}_c\cdot\text{STAT1}_c\cdot k_{11} \quad \text{(eq S46)}$$

$$\frac{d\text{ARC-STAT12}_c}{dt} = \text{ARC-STAT2}_c\cdot\text{STAT1}_c\cdot k_{11} - \text{ARC-STAT12}_c\cdot k_{13}\cdot\left(\frac{1}{1 + V_{I,N}\cdot P_{NS}}\right) - \text{ARC-STAT12}_c\cdot k_{12} \quad \text{(eq S47)}$$

$$\frac{d\text{STAT2-IRF9}_c}{dt} = \text{IRF9}_c\cdot\text{STAT2}_c\cdot k_{60} - \text{STAT2-IRF9}_c\cdot(k_{61} + k_{64}) - \text{ARC}\cdot\text{STAT2-IRF9}_c\cdot k_8$$
$$-\text{STAT2-IRF9}_c\cdot k_{36} + \text{STAT2-IRF9}_n\cdot\text{Vn2c}\cdot k_{65} \quad \text{(eq S48)}$$

$$\frac{d\text{ISGF3}_c}{dt} = \text{ISGF3-CP}\cdot k_{39} - \text{ISGF3}_c\cdot(k_{16} + k_{15}) - \text{CP}\cdot\text{ISGF3}_c\cdot k_{38} + \text{IRF9}_c\cdot\text{PSC}_c\cdot k_{14} + \text{ISGF3}_n\cdot\text{Vn2c}\cdot k_{17} \quad \text{(eq S49)}$$

$$\frac{d\text{PSC}_c}{dt} = \text{ARC-STAT12}_c\cdot k_{13}\cdot\left(\frac{1}{1 + V_{I,N}\cdot P_{\text{NS}}}\right) + \text{ISGF3}_c\cdot k_{15} - \text{PSC}_c\cdot k_{18} + \text{PSC-CP}\cdot k_{42}$$
$$-\text{CP}\cdot\text{PSC}_c\cdot k_{41} - \text{IRF9}_c\cdot\text{PSC}_c\cdot k_{14} + \text{PSC}_n\cdot\text{Vn2c}\cdot k_{19} \quad \text{(eq S50)}$$

$$\frac{d\text{ISGF3-CP}}{dt} = \text{CP}\cdot\text{ISGF3}_c\cdot k_{38} - \text{ISGF3-CP}\cdot(k_{40} + k_{39}) \quad \text{(eq S51)}$$

$$\frac{d\text{PSC-CP}}{dt} = \text{CP}\cdot\text{PSC}_c\cdot k_{41} - \text{PSC-CP}\cdot k_{43} - \text{PSC-CP}\cdot k_{42} \quad \text{(eq S52)}$$

$$\frac{d\mathrm{NP}}{dt} = \mathrm{ISGF3\text{-}NP} \cdot k_{48} + \mathrm{ISGF3\text{-}NP} \cdot k_{49} + \mathrm{B}_O\_\mathrm{NP} \cdot k_{51} + \mathrm{B}_O\_\mathrm{NP} \cdot k_{52} + \mathrm{PSC\text{-}NP} \cdot k_{45} + \mathrm{PSC\text{-}NP} \cdot k_{46} - \mathrm{ISGF3}_n \cdot \mathrm{NP} \cdot k_{47} - \mathrm{NP} \cdot \mathrm{B}_O \cdot k_{50} - \mathrm{NP} \cdot \mathrm{PSC}_n \cdot k_{44} \quad \text{(eq S53)}$$

$$\frac{d\mathrm{STAT1}_n}{dt} = \mathrm{ISGF3\text{-}NP} \cdot k_{49} + \mathrm{B}_O\_\mathrm{NP} \cdot k_{52} - \mathrm{STAT1}_n \cdot k_{57} + \mathrm{PSC\text{-}NP} \cdot k_{46} + \mathrm{STAT1}_c \cdot \mathrm{Vc2n} \cdot k_{56} \quad \text{(eq S54)}$$

$$\frac{d\mathrm{STAT2}_n}{dt} = \mathrm{ISGF3\text{-}NP} \cdot k_{49} + \mathrm{B}_O\_\mathrm{NP} \cdot k_{52} - \mathrm{STAT2}_n \cdot k_{59} + \mathrm{STAT2\text{-}IRF9}_n \cdot k_{37} + \mathrm{STAT2\text{-}IRF9}_n \cdot k_{63} + \mathrm{PSC\text{-}NP} \cdot k_{46} - \mathrm{IRF9}_n \cdot \mathrm{STAT2}_n \cdot k_{62} + \mathrm{STAT2}_c \cdot \mathrm{Vc2n} \cdot k_{58} \quad \text{(eq S55)}$$

$$\frac{d\mathrm{PIAS}}{dt} = \mathrm{PIAS\text{-}ISGF3} \cdot k_{54} - \mathrm{ISGF3}_n \cdot \mathrm{PIAS} \cdot k_{53} \quad \text{(eq S56)}$$

$$\frac{d\mathrm{PSC}_n}{dt} = \mathrm{ISGF3}_n \cdot k_{21} - \mathrm{PSC}_n \cdot k_{19} + \mathrm{PSC\text{-}NP} \cdot k_{45} - \mathrm{IRF9}_n \cdot \mathrm{PSC}_n \cdot k_{20} - \mathrm{NP} \cdot \mathrm{PSC}_n \cdot k_{44} + \mathrm{PSC}_c \cdot \mathrm{Vc2n} \cdot k_{18} \quad \text{(eq S57)}$$

$$\frac{d\mathrm{IRF9}_n}{dt} = \mathrm{ISGF3}_n \cdot k_{21} - \mathrm{IRF9}_n \cdot k_{67} - \mathrm{IRF9}_n \cdot k_{35} + \mathrm{ISGF3\text{-}NP} \cdot k_{49} + \mathrm{B}_O\_\mathrm{NP} \cdot k_{52} + \mathrm{STAT2\text{-}IRF9}_n \cdot k_{63} - \mathrm{IRF9}_n \cdot \mathrm{STAT2}_n \cdot k_{62} - \mathrm{IRF9}_n \cdot \mathrm{PSC}_n \cdot k_{20} + \mathrm{IRF9}_c \cdot \mathrm{Vc2n} \cdot k_{66} \quad \text{(eq S58)}$$

$$\frac{d\mathrm{ISGF3}_n}{dt} = \mathrm{ISGF3\text{-}NP} \cdot k_{48} - \mathrm{ISGF3}_n \cdot k_{21} - \mathrm{ISGF3}_n \cdot k_{17} + \mathrm{B}_O \cdot k_{23} + \mathrm{PIAS\text{-}ISGF3} \cdot k_{54} - \mathrm{ISGF3}_n \cdot \mathrm{NP} \cdot k_{47} - \mathrm{ISGF3}_n \cdot \mathrm{B}_U \cdot k_{22} - \mathrm{ISGF3}_n \cdot \mathrm{PIAS} \cdot k_{53} + \mathrm{IRF9}_n \cdot \mathrm{PSC}_n \cdot k_{20} + \mathrm{ISGF3}_c \cdot \mathrm{Vc2n} \cdot k_{16} \quad \text{(eq S59)}$$

$$\frac{d\mathrm{PSC\text{-}NP}}{dt} = \mathrm{NP} \cdot \mathrm{PSC}_n \cdot k_{44} - \mathrm{PSC\text{-}NP} \cdot k_{46} - \mathrm{PSC\text{-}NP} \cdot k_{45} \quad \text{(eq S60)}$$

$$\frac{d\mathrm{B}_U}{dt} = \mathrm{B}_O \cdot k_{23} + \mathrm{B}_O\_\mathrm{NP} \cdot k_{52} - \mathrm{ISGF3}_n \cdot \mathrm{B}_U \cdot k_{22} \quad \text{(eq S61)}$$

$$\frac{d\mathrm{B}_O\_\mathrm{NP}}{dt} = \mathrm{NP} \cdot \mathrm{B}_O \cdot k_{50} - \mathrm{B}_O\_\mathrm{NP} \cdot k_{52} - \mathrm{B}_O\_\mathrm{NP} \cdot k_{51} \quad \text{(eq S62)}$$

$$\frac{d\mathrm{B}_O}{dt} = \mathrm{B}_O\_\mathrm{NP} \cdot k_{51} - \mathrm{B}_O \cdot k_{23} + \mathrm{ISGF3}_n \cdot \mathrm{B}_U \cdot k_{22} - \mathrm{NP} \cdot \mathrm{B}_O \cdot k_{50} \quad \text{(eq S63)}$$

$$\frac{d\text{PIAS-ISGF3}}{dt} = \mathrm{ISGF3}_n \cdot \mathrm{PIAS} \cdot k_{53} - \text{PIAS-ISGF3} \cdot k_{54} \quad \text{(eq S64)}$$

$$\begin{aligned}\frac{d\text{STAT2-IRF9}_n}{dt} = {} & \mathrm{IRF9}_n \cdot \mathrm{STAT2}_n \cdot k_{62} - \text{STAT2-IRF9}_n \cdot k_{63} - \text{STAT2-IRF9}_n \cdot k_{65} \\ & - \text{STAT2-IRF9}_n \cdot k_{37} + \text{STAT2-IRF9}_c \cdot \mathrm{Vc2n} \cdot k_{64}\end{aligned} \quad \text{(eq S65)}$$

$$\frac{d\text{ISGF3-NP}}{dt} = \mathrm{ISGF3}_n \cdot \mathrm{NP} \cdot k_{47} - \text{ISGF3-NP} \cdot k_{49} - \text{ISGF3-NP} \cdot k_{48} \quad \text{(eq S66)}$$

$$\frac{d\mathrm{IRF7}}{dt} = k_{71} \cdot \mathrm{Vn2c} \cdot \mathrm{pIRF7} - \mathrm{IRF7} \cdot k_{\text{IRF3-IKKe-TBK1}} \cdot (\mathrm{pIKKe} + \mathrm{aTBK1}) \cdot \left(\frac{1}{1 + V_{I,N} \cdot P_{NS}}\right) + k_{79} \cdot \mathrm{IRF7\_m} - \mu_{\mathrm{IRF7}} \cdot \mathrm{IRF7} \quad \text{(eq S67)}$$

$$\frac{d\mathrm{pIRF7}}{dt} = \mathrm{IRF7} \cdot \mathrm{Vc2n} \cdot k_{\text{IRF3-IKKe-TBK1}} \cdot (\mathrm{pIKKe} + \mathrm{aTBK1}) \cdot \left(\frac{1}{1 + V_{I,N} \cdot P_{NS}}\right) - k_{71} \cdot \mathrm{pIRF7} \quad \text{(eq S68)}$$

$$\frac{d\mathrm{ISGav\_m}}{dt} = \mathrm{B}_O \cdot \mathrm{Vn2c} \cdot k_{\mathrm{ISGav\_m}} - \mathrm{ISGav\_m} \cdot \mu_{\text{ISG RNA}} + \mathrm{Vn2c} \cdot k_{\mathrm{m\_ISGav}} \cdot \mathrm{pIRF3} \quad \text{(eq S69)}$$

$$\frac{d\mathrm{ISGav}}{dt} = \mathrm{ISGav\_m} \cdot k_{\text{t, ISG RNA}} - \mathrm{ISGav} \cdot \mu_{\mathrm{ISGav}} \quad \text{(eq S70)}$$

$$\frac{d\mathrm{ISGn\_m}_n}{dt} = k_{72} \cdot \mathrm{B}_O - k_{73} \cdot \mathrm{ISGn\_m}_n \quad \text{(eq S71)}$$

$$\frac{d\mathrm{IRF9\_m}_n}{dt} = k_{74} \cdot \mathrm{B}_O - k_{75} \cdot \mathrm{IRF9\_m}_n \quad \text{(eq S72)}$$

$$\frac{d\mathrm{IRF7\_m}}{dt} = k_{76} \cdot \mathrm{B}_O - \left(\frac{\log_e(2)}{\tau_5}\right) \cdot \mathrm{IRF7\_m} \quad \text{(eq S73)}$$

$$\frac{d\text{ISGn\_m}_c}{dt} = \text{Vn2c} \cdot \text{ISGn\_m}_n \cdot k_{73} - k_{77} \cdot \text{ISGn\_m}_c \qquad \text{(eq S74)}$$

$$\frac{d\text{IRF9\_m}_c}{dt} = \text{Vn2c} \cdot \text{IRF9\_m}_n \cdot k_{75} - k_{78} \cdot \text{IRF9\_m}_c \qquad \text{(eq S75)}$$

### Supplementary Tables

Integrative Modelling of Innate Immune Response Dynamics during Virus Infection

Ramya Boddepalli, Harsh Chhajer, and Rahul Roy

| Eqn no | Species | Description | Value | Ref. |
|---|---|---|---|---|
| S1 | $V_0$ | Virus Input | 100 nM | High MOI |
| S2 | $V_T$ | Viral Titre | 0 | |
| S3 | $V_I$ | Internalized virus | 0 | |
| S4 | $R_{cyt}$ | Cytoplasmic viral RNA | 0 | |
| S5 | $R_{CM}$ | (+) RNA in CMs | 0 | |
| S6 | $P_S$ | Viral Structural Proteins | 0 | |
| S7 | $P_{NS}$ | Viral Non-Structural Proteins | 0 | |
| S8 | $RC_{CM}$ | Compartmentalized replication complexes | 0 | |
| S9 | $RNA_{ds}$ | dsRNA in cytoplasm | 0 | |
| S10 | RIGI | PRR sensor | 5.34 nM | [1] |
| S11 | aRIGI | Activated RIGI | 0 | |
| S12 | MAVS | Mitochondrial antiviral-signaling protein | 277.76 nM | [1] |
| S13 | aMAVS | Activated MAVS | 0 | |
| S14 | IKKe | I$\kappa$B kinase $\varepsilon$ | 3.08 nM | [1] |
| S15 | aIKKe | Activated IKKe | 0 | |
| S16 | TBK1 | TANK-binding kinase 1 | 97.17 nM | [1] |
| S17 | aTBK1 | Activated TBK1 | 0 | |
| S18 | IRF3 | Interferon regulatory factor 3 | 37.86 nM | [1] |
| S19 | pIRF3 | Phosphorylated IRF3 | 0 | |
| S20 | IKK | Inhibitor of kappa B kinase | 37.97 nM | [1] |
| S21 | aIKK | Activated IKK | 0 | |
| S22 | NFkB-IkBa$_c$ | Inactive NF$\kappa$B in cytoplasm | 11.36 nM | [1] |
| S23 | pNFkB$_n$ | Activated NF$\kappa$B in nucleus | 0 | |
| S24 | NFkB$_n$ | NF$\kappa$B in nucleus | 0 | |
| S25 | NFkB$_c$ | NF$\kappa$B in cytoplasm | 101.73 nM | [1] |
| S26 | IkBa$_c$ | inhibitor of kappa B in cytoplasm | 0 | |
| S27 | IFNb_m | Interferon $\beta$ mRNA | 0 | |
| S28 | IFNa_m | Interferon $\alpha$ mRNA | 0 | |
| S29 | IFNl_m | Interferon $\lambda$ mRNA | 0 | |
| S30 | $IFN_c$ | Interferon in cytoplasm | | |
| S31 | $IFNl_c$ | Interferon $\lambda$ in cytoplasm | 0 | |
| S32 | JAK | Janus Kinase 1 | 151.86 nM | [1] |
| S33 | RJC | Receptor JAK complex | 0 | |
| S34 | $STAT1_c$ | STAT 1 in cytoplasm | 1114.68 nM | [1] |
| S35 | CP | Cytoplasmic Phosphatase | 20 nM | [2] |
| S36 | ISGn | Negative regulator ISG | 0 | |
| S37 | IFNex | Extracellular interferon type-1 | 0 | |
| S38 | $STAT2_c$ | STAT2 in cytoplasm | 6.50 nM | [1] |
| S39 | TYK | Tyrosine Kinase 2 | 20.70 nM | [1] |
| S40 | ARC | Activated Receptor Complex | 0 | |
| S41 | RTC | Receptor TYK Complex | 0 | |

Table S1: **Description of model variables and their initial condition values used in simulation**

| Eqn no | Species | Description | Value | Ref. |
|---|---|---|---|---|
| S42 | IFNAR1 | Interferon alpha receptor subunit 1 | 1000 nM | [2] |
| S43 | IFNAR2 | Interferon alpha receptor subunit 2 | 1000 nM | [2] |
| S44 | IFNARd | Interferon receptor dimer | 0 | |
| S45 | $IRF9_c$ | IRF9 in cytoplasm | 45 nM | [2] |
| S46 | $ARC\text{-}STAT2_c$ | ARC and STAT2 complex in cytoplasm | 0 | |
| S47 | $ARC\text{-}STAT12_c$ | ARC, STAT2 and STAT1 complex in cytoplasm | 0 | |
| S48 | STAT2-IRF9c | STAT2 and IRF9 complex in cytoplasm | 0 | |
| S49 | $ISGF3_c$ | ISGF3 in cytoplasm | 0 | |
| S50 | $PSC_c$ | Phosphorylated STATs complex in cytoplasm | 0 | |
| S51 | ISGF3-CP | ISGF3 and CP complex | 0 | |
| S52 | PSC-CP | PSC and CP complex | 0 | |
| S53 | NP | Nuclear Phosphatase | 40 nM | [2] |
| S54 | $STAT1_n$ | STAT1 in nucleus | 0 | |
| S55 | $STAT2_n$ | STAT2 in nucleus | 0 | |
| S56 | PIAS | Protein inhibitor of activated STAT | 41.96 nM | [1] |
| S57 | $PSC_n$ | Phosphorylated STATs complex in nucleus | 0 | |
| S58 | $IRF9_n$ | IRF9 in nucleus | 0 | |
| S59 | $ISGF3_n$ | ISGF3 in nucleus | 0 | |
| S60 | PSC-NP | PSC and NP complex | 0 | |
| S61 | $B_U$ | Open ISGF3 binding sites | 500 nM | [2] |
| S62 | $B_O$-NP | Occupied ISGF3 binding sites—NP | 0 | |
| S63 | $B_O$ | Occupied ISGF3 binding sites | 0 | |
| S64 | PIAS-ISGF3 | PIAS and ISGF3 complex | 0 | |
| S65 | $STAT2\text{-}IRF9_n$ | STAT2 and IRF9 complex in nucleus | 0 | |
| S66 | ISGF3-NP | ISGF3 and NP complex | 0 | |
| S67 | IRF7 | Interferon regulatory factor 7 | 24 nM | [3] |
| S68 | pIRF7 | Phosphorylated IRF7 | 0 | |
| S69 | ISGav_m | Antiviral ISG mRNA | 0 | |
| S70 | ISGav | Antiviral ISG | 0 | |
| S71 | $ISGn\_m_n$ | Negative regulator ISG mRNA in nucleus | 0 | |
| S72 | $IRF9\_m_n$ | IRF9 mRNA in nucleus | 0 | |
| S73 | IRF7_m | IRF7 mRNA | 0 | |
| S74 | $ISGn\_m_c$ | Negative regulator ISG mRNA in cytoplasm | 0 | |
| S75 | $IRF9\_m_c$ | IRF9 mRNA in cytoplasm | 0 | |

Table S2: **Description of model variables and their initial condition values used in simulation (contd..)**

| Parameter | Description | Value | Ref. |
|---|---|---|---|
| $k_{\mathrm{en,V}}$ | Rate of virus entry | 0.43 $h^{-1}$ | [4] |
| $k_{f,V}$ | Rate of fusion of the viral and endosomal membranes | 0.031 $h^{-1}$ | [4] |
| $\mu_{V_I}$ | Decay rate of internalised virus | 0.04 $h^{-1}$ | [4] |
| $k_{a,V}$ | Effective virus generation rate | $3.6\times10^{-8}$ molecules$^{-1}$ $h^{-1}$ | [5] |
| $k_{e,V}$ | Export rate of $R_{CM}$ into the cytoplasm | $6.6\times10^{-2}$ $h^{-1}$ | [5] |
| $k_{r,V}$ | (+)RNA synthesis rate per $RC_{CM}$ | 3.6 $h^{-1}$ | [5] |
| $k_{c,V}$ | Formation rate of $RC_{CM}$ | $2.6\times10^{-3}$ molecules$^{-1}$ $h^{-1}$ | [5] |
| $k_{t,V}$ | Protein production rate per $R_{cyt}$ | 23.7 $h^{-1}$ | [5] |
| $\tau$ | Time constant for functional development of CMs | 5.8 h | [5] |
| $k_{l,V}$ | Rate of transfer of dsRNA into cytoplasm | 0.028 $h^{-1}$ | [4] |
| $N_{\mathrm{C,\,V}}$ | $RC_{CM}$ carrying capacity of the host cell | 88 | [5] |
| $\mu_{r,V}$ | Degradation rate of $R_{cyt}$ | 0.25 $h^{-1}$ | [5] |
| $\mu_{p,V}$ | Degradation rate of PS and PNS | 0.11 $h^{-1}$ | [5] |
| $\mu_{V,V}$ | Degradation rate of extracellular virus, $V_T$ | $6\times10^{-3}$ $h^{-1}$ | [5] |
| nSP | Number of PS per virus particle | 180 | [5] |
| $k_{69}$ | Dissociation rate of ARC by ISGn (negative regulator ISG) | 0.203 $nM^{-1}$ $min^{-1}$ | [6] |
| $k_{70}$ | Translation rate of IRF9 mRNA | 0.864 $min^{-1}$ | [6] |
| $k_{71}$ | Rate of dephosphorylation of pIRF7 | $10^{-3}$ $min^{-1}$ | [1] |
| $k_{72}$ | Rate of production of ISGn mRNA | 0.147 $min^{-1}$ | [6] |
| $k_{73}$ | Translocation of ISGn mRNA from nucleus to cytoplasm | 4.89 $min^{-1}$ | [6] |
| $k_{74}$ | Rate of production of IRF9 mRNA | 0.33 $min^{-1}$ | [6] |
| $k_{75}$ | Translocation rate of IRF9 mRNA from cytoplasm to nucleus | 0.876 $min^{-1}$ | [6] |
| $k_{76}$ | Rate of production of IRF7 mRNA | $6\times10^{-5}$ $min^{-1}$ | [6] |
| $k_{77}$ | Degradation rate of ISGn mRNA | $4.36\times10^{-2}$ $min^{-1}$ | [6] |
| $k_{78}$ | Degradation rate of IRF9 mRNA | $1.39\times10^{-2}$ $min^{-1}$ | [6] |
| $k_{79}$ | Translation rate of IRF7 mRNA | 43.87 $min^{-1}$ | [7] |
| $k_{transISGn}$ | Translation rate of IRF9 mRNA | 0.63 $min^{-1}$ | [6] |
| kinhISGn | Inhibition of formation of ARC by ISGn (negative regulator ISG) | 889.4 $nM^{-1}$ | [8] |

Table S3: **Description of model parameters and their values (HCV) used unless specified otherwise**

| Parameter | Description | Value | Ref. |
|---|---|---|---|
| $k_{a,V}$ | Effective virus generation rate | $8\times10^{-9}$ molecules$^{-1}$ h$^{-1}$ | [5] |
| $k_{e,V}$ | Export rate of $R_{CM}$ into the cytoplasm | $7.2\times10^{-2}$ h$^{-1}$ | [5] |
| $k_{r,V}$ | (+)RNA synthesis rate per $RC_{CM}$ | 3.7 h$^{-1}$ | [5] |
| $k_{c,V}$ | Formation rate of $RC_{CM}$ | $1.6\times10^{-2}$ molecules$^{-1}$ h$^{-1}$ | [5] |
| $k_{t,V}$ | Protein production rate per $R_{cyt}$ | $1.6\times10^{2}$ h$^{-1}$ | [5] |
| $\tau$ | Time constant for functional development of CMs | 2.6 h | [5] |
| $N_{\text{C, V}}$ | $RC_{CM}$ carrying capacity of the host cell | $1.21\times10^{3}$ | [5] |
| $\mu_{r,V}$ | Degradation rate of $R_{cyt}$ | 0.25 h$^{-1}$ | [5] |
| $\mu_{p,V}$ | Degradation rate of PS and PNS | 0.11 h$^{-1}$ | [5] |
| $\mu_{V,V}$ | Degradation rate of extracellular virus, $V_T$ | $6\times10^{-3}$ h$^{-1}$ | [5] |
| nSP | Number of PS per virus particle | 180 | [5] |

Table S4: **Description of model parameters and their values for JEV**

| Parameter | Description | Value | Ref. |
|---|---|---|---|
| $\mu_{IRF7}$ | Degradation rate of IRF7 | $1.67\times10^{-3}$ min$^{-1}$ | [9] |
| degARCISGn | Degradation rate of ARC by ISGn (negative regulator ISG) | 0.0147 nM$^{-1}$ min$^{-1}$ | [8] |
| degRecISGn | Degradation rate of Receptor by ISGn (negative regulator ISG) | $1.7\times10^{-4}$ nM$^{-1}$ min$^{-1}$ | [8] |
| $k_s$ | Rate of IFN secretion | $1.65\times10^{-2}$ min$^{-1}$ | [4] |
| $\tau_5$ | Half-life of IRF7 mRNA | 120 min | [10] |
| $k_{m\_IFNa}$ | IFN $\alpha$ mRNA production rate | $1.67\times10^{-4}$ nM$^{-1}$ min$^{-1}$ | [1] |
| $k_{m\_IFNb}$ | IFN $\beta$ mRNA production rate | $1.99\times10^{-4}$ nM$^{-1}$ min$^{-1}$ | [1] |
| $k_{m\_IFNl}$ | IFN $\lambda$ mRNA production rate | 0.154 min$^{-1}$ | [1] |
| $k_{m\_ISGav}$ | IFN-independent antiviral ISG mRNA production rate | $7\times10^{-3}$ min$^{-1}$ | [1] |
| $k_{ISGav\_m}$ | IFN-dependent antiviral ISG mRNA production rate | 15.41 min$^{-1}$ | [1] |
| $\mu_{m\_IFNa}$ | Degradation rate of IFN $\alpha$ mRNA | $2.3\times10^{-3}$ min$^{-1}$ | [1] |
| $\mu_{m\_IFNb}$ | Degradation rate of IFN $\beta$ mRNA | $2.8\times10^{-3}$ min$^{-1}$ | [1] |
| $\mu_{m\_IFNl}$ | Degradation rate of IFN $\lambda$ mRNA | $3.6\times10^{-3}$ min$^{-1}$ | [1] |
| $\mu_{ISGRNA}$ | Degradation rate of antiviral ISG mRNA | $2.8\times10^{-3}$ min$^{-1}$ | [1] |
| $\mu_{ISGav}$ | Degradation rate of antiviral ISG | $2.1\times10^{-4}$ min$^{-1}$ | [1] |
| $V_{\mathrm{cyt}}$ | Cytoplasmic volume | $1.2\times10^{-12}$ L | [1] |
| $V_{\mathrm{n}}$ | Nuclear volume | $4.66\times10^{-13}$ L | [1] |
| Vc2n | Volume ratio: cytoplasm to nucleus | 2.585 | – |
| Vn2c | Volume ratio: nucleus to cytoplasm | 0.387 | – |
| $b_{\mathrm{IRF3}}$ | Dephosphorylation rate of IRF3 | $10^{-3}$ min$^{-1}$ | [1] |
| $b_{\mathrm{MAVS}}$ | Deactivation rate of MAVS | 0.219 min$^{-1}$ | [3] |
| $b_{\mathrm{RIGI}}$ | Deactivation rate of RIGI | $9.3\times10^{-2}$ min$^{-1}$ | [1] |
| $b_{\mathrm{KINASE}}$ | Deactivation rate of TBK1, IKK$\epsilon$ and IKK | $8\times10^{-2}$ min$^{-1}$ | [1] |
| $k_{\mathrm{trans\_IFNl}}$ | Translation rate of IFN $\lambda$ | $6.2\times10^{-4}$ min$^{-1}$ | [1] |
| $k_{\mathrm{transp\_NF\kappa B}}$ | Translocation rate of NF$\kappa$B from nucleus to cytoplasm | $2.3\times10^{-4}$ min$^{-1}$ | [1] |
| $\mu_{IFN}$ | degradation rate of IFN | $2.5\times10^{-3}$ min$^{-1}$ | [4] |
| $\mu_{IFNl}$ | degradation rate of IFN$\lambda$ | $1.6\ \times10^{-4}$ min$^{-1}$ | [1] |
| $\mu_{I\kappa B\alpha}$ | degradation rate of I$\kappa$B$\alpha$ | $6.8\ \times10^{-3}$ min$^{-1}$ | [1] |
| $\mu_{RIGI}$ | degradation rate of RIGI | $1.4\ \times10^{-3}$ min$^{-1}$ | [1] |
| $k_{\mathrm{t,\ ISG\ RNA}}$ | translation rate of antiviral ISG (ISGav) | $1.2\times10^{-4}$ min$^{-1}$ | [1] |

Table S5: **Description of model parameters and their values used unless specified otherwise**

| Parameter | Description | Value | Ref. |
|---|---|---|---|
| $k_1$ | TYK and IFNAR1 association rate | 6 $nM^{-1}$ $min^{-1}$ | [2] |
| $k_{10}$ | Dissociation rate of ARC and $STAT2_c$ from ARC-STAT$2_c$ | 240 $min^{-1}$ | [2] |
| $k_{11}$ | Association rate of $STAT1_c$ with $ARC - STAT2_c$ | 0.12 $nM^{-1}$ $min^{-1}$ | [2] |
| $k_{12}$ | Dissociation rate of $STAT1_c$ from $ARC - STAT2_c$ | 240 $min^{-1}$ | [2] |
| $k_{13}$ | Formation of $PSC_c$ | 480 $min^{-1}$ | [2] |
| $k_{14}$ | Association rate of $PSC_c$ with $IRF9_c$ resulting in $ISGF3_c$ | 6 $nM^{-1}$ $min^{-1}$ | [2] |
| $k_{15}$ | Disassociation rate of $ISGF3_c$ into $PSC_c$ and $IRF9_c$ | 6 $min^{-1}$ | [2] |
| $k_{16}$ | Translocation rate of $ISGF3_c$ to nucleus | 0.9 L $min^{-1}$ | [2] |
| $k_{17}$ | Translocation rate of $ISGF3_n$ to cytoplasm | 0 L $min^{-1}$ | [2] |
| $k_{18}$ | Translocation rate of $PSC_c$ to nucleus | 0.9 L $min^{-1}$ | [2] |
| $k_{19}$ | Translocation rate of $PSC_n$ to cytoplasm | 0 L $min^{-1}$ | [2] |
| $k_2$ | TYK and IFNAR1 disassociation rate | 3 $min^{-1}$ | [2] |
| $k_{20}$ | Association rate of $PSC_n$ with $IRF9_n$ resulting in $ISGF3_n$ | 0.6 $nM^{-1}$ $min^{-1}$ | [2] |
| $k_{21}$ | Dissociation rate of $ISGF3_n$ into $IRF9_n$ and $PSC_n$ | 0.6 $min^{-1}$ | [2] |
| $k_{22}$ | Association rate of $ISGF3_n$ to the $B_U$ | 6 $nM^{-1}$ $min^{-1}$ | [2] |
| $k_{23}$ | Dissociation rate of $B_O$ into $ISGF3_n$ and $B_U$ | 6 $min^{-1}$ | [2] |
| $k_{27}$ | Basal production of IRF$9_c$ | 0.3 nM $min^{-1}$ | [2] |
| $k_{29}$ | Degradation rate of IRF$9_c$ | 0.006 $min^{-1}$ | [2] |
| $k_3$ | JAK and IFNAR2 association rate | 6 $nM^{-1}$ $min^{-1}$ | [2] |
| $k_{31}$ | Degradation rate of ISGn | 0.03 $min^{-1}$ | [2] |
| $k_{32}$ | Dissociation rate of ARC | 0.018 $min^{-1}$ | [2] |
| $k_{34}$ | Deactivation rate of ARC into IFNARd | 0.6 $min^{-1}$ | [2] |
| $k_{35}$ | Degradation rate of $IRF9_n$ | 0.006 $min^{-1}$ | [2] |
| $k_{36}$ | Dissociation of $STAT2$-$IRF9_c$ into $STAT2_c$ | 0.006 $min^{-1}$ | [2] |
| $k_{37}$ | Dissociation of $STAT2$-$IRF9_n$ into $STAT2_n$ | 0.006 $min^{-1}$ | [2] |
| $k_{38}$ | Association of $ISGF3_c$ with CP | 0.06 $nM^{-1}$ $min^{-1}$ | [2] |
| $k_{39}$ | Dissociation of ISGF3-CP into $ISGF3_c$ and CP | 12 $min^{-1}$ | [2] |
| $k_4$ | JAK and IFNAR2 disassociation rate | 3 $min^{-1}$ | [2] |
| $k_{40}$ | Dephosphorylation of $ISGF3_c$ complex by CP | 0.18 $min^{-1}$ | [2] |
| $k_{41}$ | Association of $PSC_c$ with CP | 0.06 $nM^{-1}$ $min^{-1}$ | [2] |
| $k_{42}$ | Dissociation of PSC-CP into $PSC_c$ and CP | 12 $min^{-1}$ | [2] |

Table S6: **Parameter values. Notation has been kept similar to the references mentioned.**

| Parameter | Description | Value | Ref. |
|---|---|---|---|
| $k_{43}$ | Dephosphorylation of $PSC_c$ complex by CP | 0.18 $min^{-1}$ | [2] |
| $k_{44}$ | Association of $PSC_n$ with NP | 0.6 $nM^{-1}$ $min^{-1}$ | [2] |
| $k_{45}$ | Dissociation of PSC-NP into $PSC_n$ and NP | 6 $min^{-1}$ | [2] |
| $k_{46}$ | Dephosphorylation of $PSC_n$ complex by NP | 6 $min^{-1}$ | [2] |
| $k_{47}$ | Association of $ISGF3_n$ with NP | 0.6 $nM^{-1}$ $min^{-1}$ | [2] |
| $k_{48}$ | Dissociation of ISGF3-NP into $ISGF3_n$ and NP | 6 $min^{-1}$ | [2] |
| $k_{49}$ | Dephosphorylation of $ISGF3_n$ complex by NP | 0.12 $min^{-1}$ | [2] |
| $k_5$ | Association rate of free IFN with RJC and RTC resulting in IFNARd | 0.6 $nM^{-2}$ $min^{-1}$ | [2] |
| $k_{50}$ | Association of $B_O$ with NP | 0.006 $nM^{-1}$ $min^{-1}$ | [2] |
| $k_{51}$ | Dissociation of $B_O$-$NP$ into $B_O$ and NP | 6 $min^{-1}$ | [2] |
| $k_{52}$ | Dissociation of $B_O$ into subunits | 6 $min^{-1}$ | [2] |
| $k_{53}$ | Association of $ISGF3_n$ with PIAS | 6 $nM^{-1}$ $min^{-1}$ | [2] |
| $k_{54}$ | Dissociation of ISGF3-PIAS into $ISGF3_n$ and PIAS | 6 $min^{-1}$ | [2] |
| $k_{56}$ | Translocation rate of $STAT1_c$ to nucleus | 0.075 $nM^{-1}$ $min^{-1}$ | [2] |
| $k_{57}$ | Translocation rate of $STAT1_n$ to cytoplasm | 0.6 L $min^{-1}$ | [2] |
| $k_{58}$ | Translocation rate of $STAT2_c$ to nucleus | 0.0049 L $min^{-1}$ | [2] |
| $k_{59}$ | Translocation rate of $STAT2_n$ to cytoplasm | 0.084 L $min^{-1}$ | [2] |
| $k_6$ | Disassociation rate of free IFN from RJC and RTC | 0.6 $min^{-1}$ | [2] |
| $k_{60}$ | Association of $STAT2_c$ and $IRF9_c$ | 0.6 $nM^{-1}$ $min^{-1}$ | [2] |
| $k_{61}$ | Dissociation of $STAT2-IRF9_c$ into $STAT2_c$ $IRF9_c$ | 0.6 $min^{-1}$ | [2] |
| $k_{62}$ | Association of $STAT2_n$ and $IRF9_n$ | 0.6 $nM^{-1}$ $min^{-1}$ | [2] |
| $k_{63}$ | Dissociation of $STAT2-IRF9_n$ into $STAT2_n$ $IRF9_n$ | 0.6 $min^{-1}$ | [2] |
| $k_{64}$ | Translocation rate of $STAT2-IRF9_c$ to nucleus | 0.075 L $min^{-1}$ | [2] |
| $k_{65}$ | Translocation rate of $STAT2-IRF9_n$ to cytoplasm | 0.084 L $min^{-1}$ | [2] |
| $k_{66}$ | Translocation rate of $IRF9_c$ to nucleus | 1.2 L $min^{-1}$ | [2] |
| $k_{67}$ | Translocation rate of $IRF9_n$ to cytoplasm | 0.3 L $min^{-1}$ | [2] |
| $k_7$ | Formation of ARC from IFNARd | 0.3 $min^{-1}$ | [2] |

Table S7: **Parameter values. Notation has been kept similar to the references mentioned.**

| Parameter | Description | Value | Ref. |
|---|---|---|---|
| $k_8$ | Association of STAT2c with ARC from STAT2c$_{I}RF9$ | 0.12 $nM^{-1}$ $min^{-1}$ | [2] |
| $k_9$ | Association of free STAT2c with ARC | 0.12 $nM^{-1}$ $min^{-1}$ | [2] |
| $k_{IFN}$ | Translation rate of IFN mRNA | 5.98 $\times 10^{-5}$ $min^{-1}$ | [1] |
| $k_{IKK}$ | Activation of IKK | $4.82\times 10^{-2}$ $nM^{-1}$ $min^{-1}$ | [1] |
| $k_{IKKe\text{-}TBK1}$ | Activation of IKK$\epsilon$ and TBK1 | $10^{-3}$ $nM^{-1}$ $min^{-1}$ | [1] |
| $k_{IRF3\text{-}IKKe\text{-}TBK1}$ | IRF3 phosphorylation rate by IKK$\epsilon$ and TBK1 | 0.0170 $nM^{-1}$ $min^{-1}$ | [1] |
| $k_{MAVS}$ | Activation of MAVS | $9\times 10^{-3}$ $nM^{-1}$ $min^{-1}$ | [3] |
| $k_{RIGI}$ | Activation of RIGI | 0.01 $nM^{-1}$ $min^{-1}$ | [3] |
| $k_{TFBS\_IFN\alpha}$ | IFN$\alpha$ induction by JAK-STAT pathway | 0 $min^{-1}$ | [1] |
| $k_{TFBS\_IFN\beta}$ | IFN $\beta$ induction by JAK-STAT pathway | 0 $min^{-1}$ | [1] |
| $k_{TFBS\_IFN\lambda}$ | IFN$\lambda$ induction by JAK-STAT pathway | 0 $min^{-1}$ | [1] |
| $k_{act}$ | Activation of inactive NF$\kappa$B by IKK | 0.00211 $nM^{-1}$ $min^{-1}$ | [1] |
| $k_{deph}$ | Dephosphorylation rate constant of nuclear pNF$\kappa$B | 0.0519 $min^{-1}$ | [1] |
| $k_{expr,\ I\kappa B\alpha}$ | I$\kappa$B$\alpha$ production rate | $9.2\times 10^{-5}$ $min^{-1}$ | [1] |
| $k_{inh\_p65}$ | Sequestering of NF$\kappa$B by I$\kappa$B$\alpha$ | 0.03 $nM^{-1}$ $min^{-1}$ | [1] |
| $k_{RIGI\_synt}$ | Basal level RIGI synthesis | 7.5 $\times 10^{-3}$ nM $min^{-1}$ | [1] |

Table S8: **Parameter values. Notation has been kept similar to the references mentioned.**

### Supplementary Figures

Integrative Modelling of Innate Immune Response Dynamics during Virus Infection

Ramya Boddepalli, Harsh Chhajer, and Rahul Roy

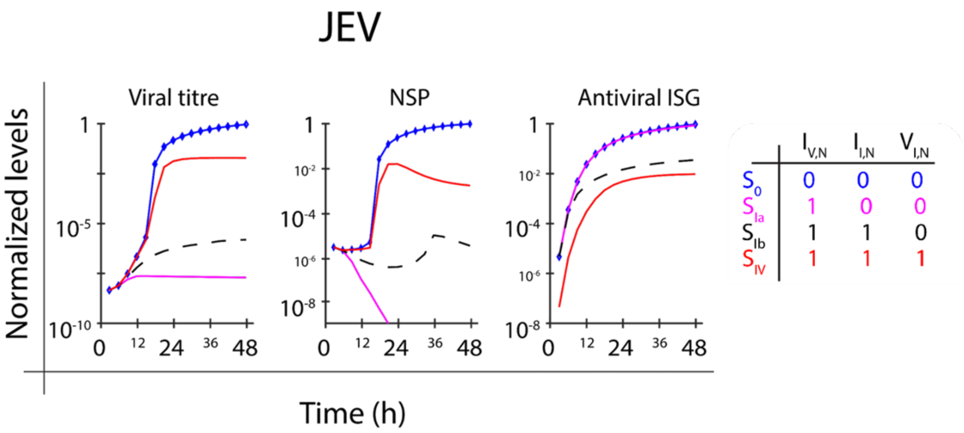


Figure S1: **Integrated mathematical model of the viral life cycle with interferon pathways** Normalised dynamics (for JEV) of the levels of viral titre (left), non-structural proteins (center), and antiviral ISGs (right) are shown and compared for different combinations of strengths of immune self-regulation ($I_{I,N}$), viral countermeasures ($V_{I,N}$) and antiviral immune response ($I_{V,N}$). The blue line represents a no-interaction ($S_0$) scenario where $I_{V,N}$, $I_{I,N}$, $V_{I,N}$ are all set to zero, whereas the red solid line represents a fully interacting system where all three are set to one ($S_{IV}$). The black dashed and magenta lines correspond to intermediate interactions ($S_{Ib}$) where '$I_{V,N} = I_{I,N} = 1$ and $V_{I,N} = 0$' and scenario ($S_{Ia}$) '$I_{V,N} = 1$ $I_{I,N} = V_{I,N} = 0$' respectively.

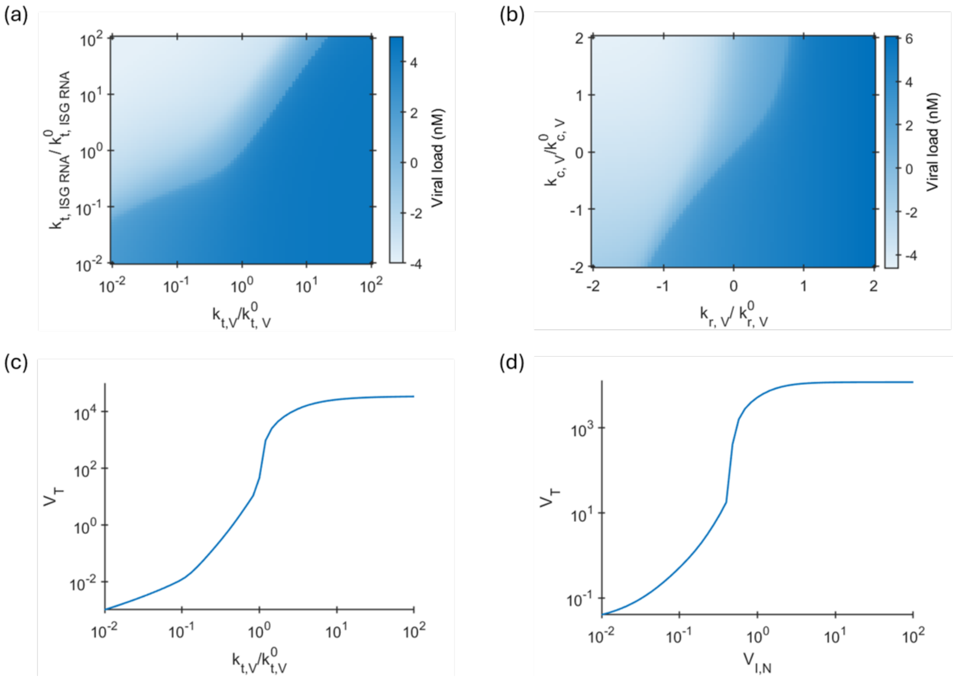


Figure S2: **Bifurcation in system dynamics**
The heatmap of the viral titer output ($V_T$) during HCV infection is shown when $k_{t,V}$ and $k_{t,ISGRNA}$ (left) and $k_{c,V}$ and $k_{r,V}$ (right). Darker shades indicate higher viral load, highlighting the nonlinear response of viral output to changes in host-virus interaction parameters.

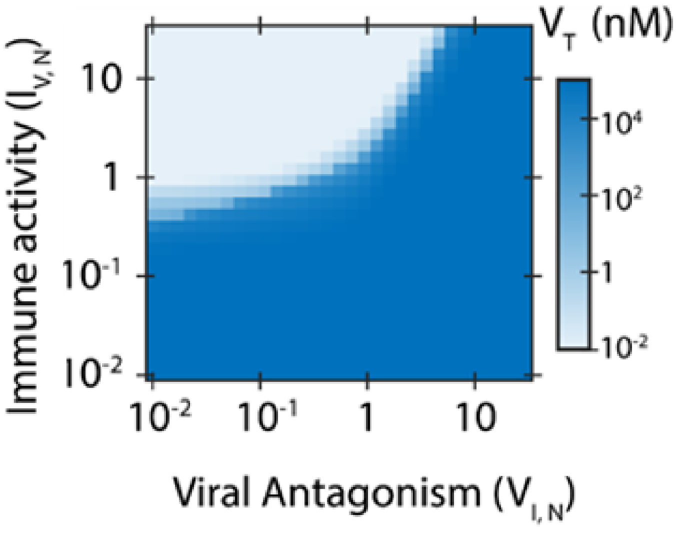


Figure S3: **Bifurcation in system dynamics**
The heatmap of the viral titer output ($V_T$) during JEV infection is shown as a function of $V_{I,N}$ and immune action strength ($I_{V,N}$). Darker shades indicate higher viral load, highlighting the nonlinear response of viral output to changes in host-virus interaction parameters.

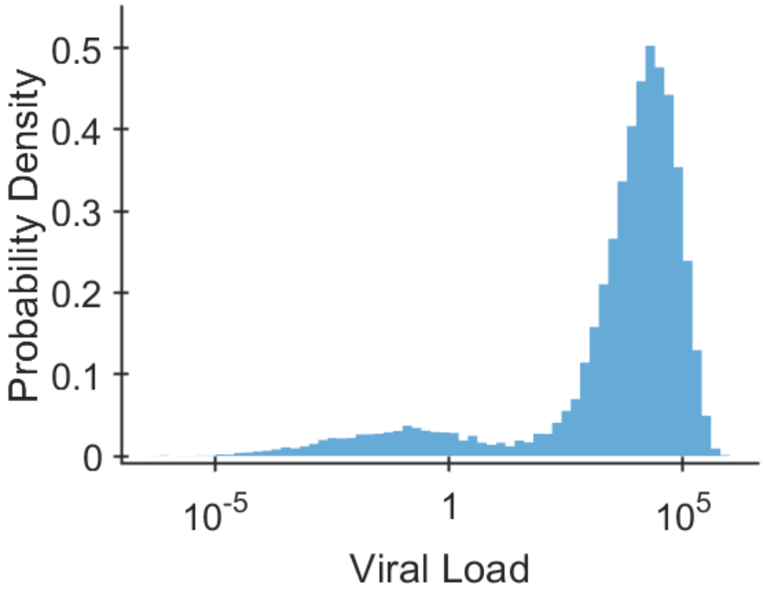


Figure S4: Distribution of Hepatitis C viral load predicted from 25,000 parameter sets generated by Latin Hypercube Sampling (LHS), with all model parameters varied within ±10% of their nominal values (90–110%). Here we fix $V_{I,N} = 0.44$ and $I_{V,N} = 1$ (critical point).

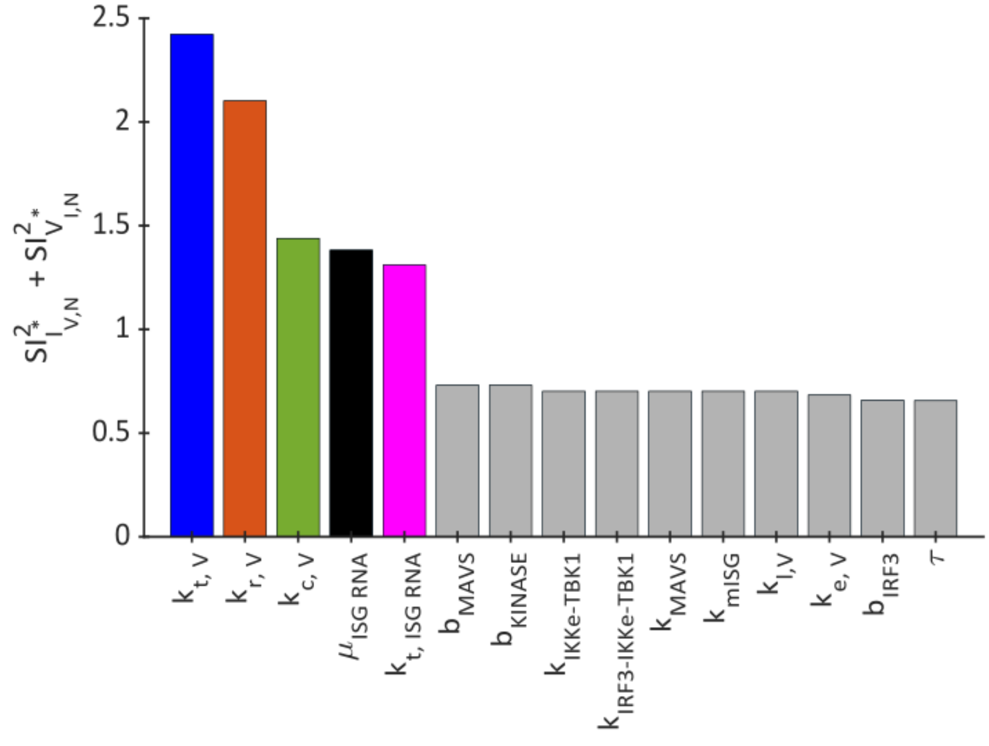


Figure S5: Bar plot showing the parameter sum of sensitivity indices of critical values (corresponding to bifurcation) of $I^*_{V,N}$ and $V^*_{I,N}$. Colored bars denote the most influential parameters: $k_{t,V}$, $k_{r,V}$, $k_{c,V}$, $\mu_{r,ISGRNA}$ and $k_{t,ISGRNA}$ (Description in Supplementary Table S3-S7).

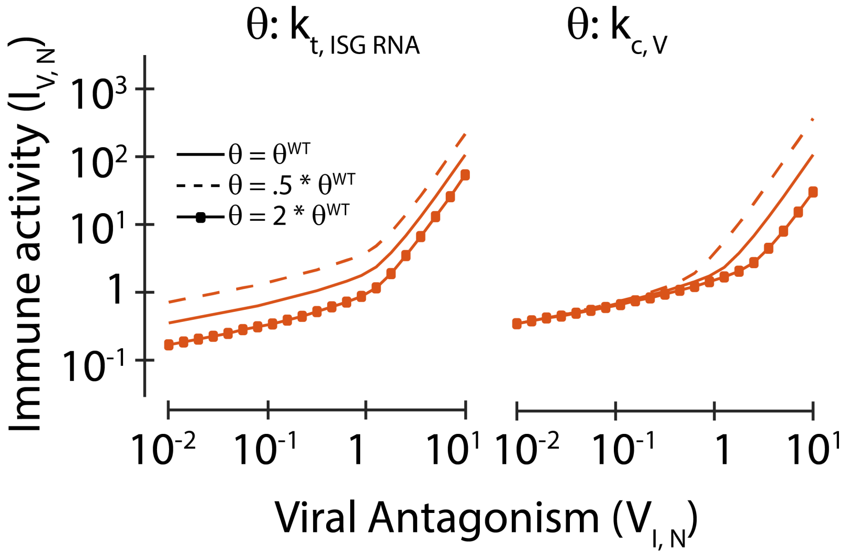


Figure S6: The bifurcation boundary for JEV infection systems are shown for different values of $k_{t,ISGRNA}$ (translation rate of ISG mRNA, left) and $k_{c,V}$ (compartmentalization rate of viral dsRNA, right). The solid line, dashed line, and the solid line with square symbols show the bifurcation boundary when the value of the parameter of interest is unperturbed, reduced by half, and doubled, respectively.

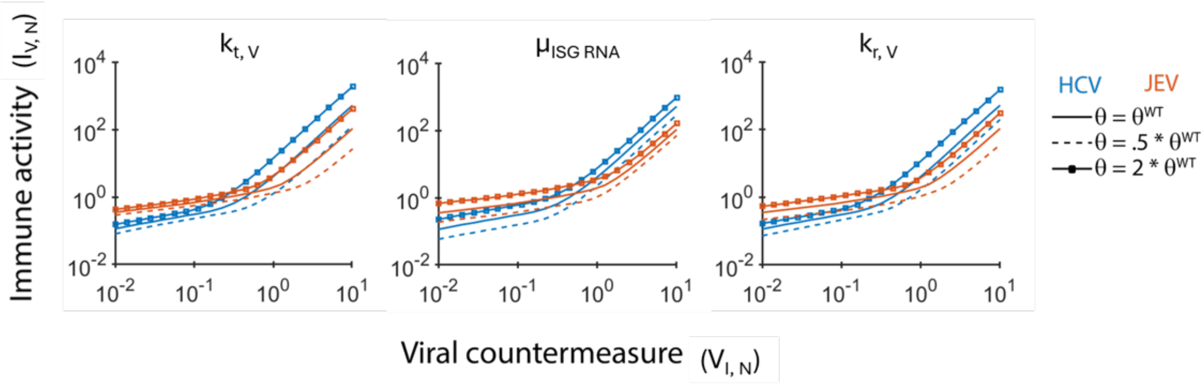


Figure S7: The bifurcation fronts for HCV (blue) and JEV (orange) infection systems are shown for different values of $k_{t,V}$ (translation rate of viral RNA, left), $\mu_{ISGRNA}$ (degradation of ISG mRNA, middle) and $k_{r,V}$ (transcription rate of viral RNA, right). The solid line, dashed line, and the solid line with square symbols show the bifurcation boundary when the value of the parameter of interest is unperturbed, reduced by half, and doubled, respectively.

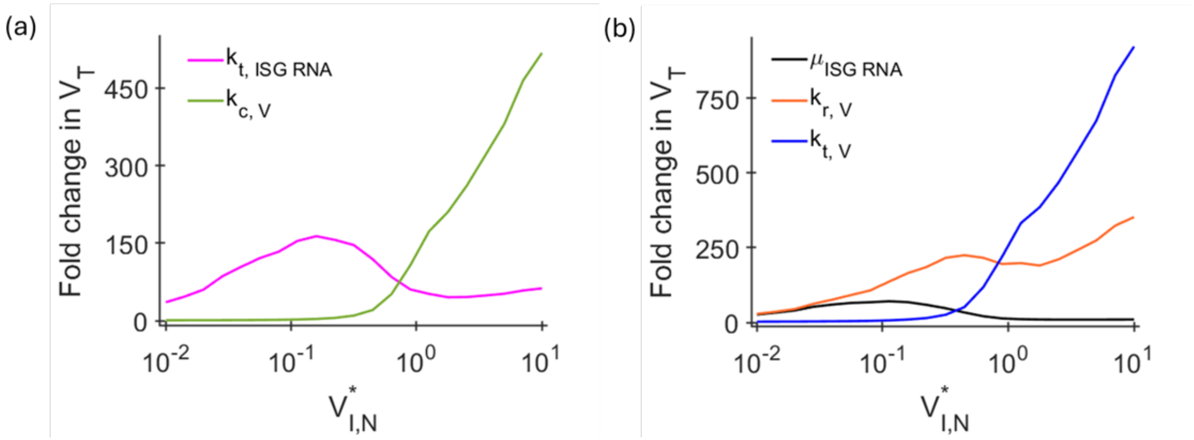


Figure S8: The line plots show the fold change in the steady state values of JEV $V_T$ at points along the front when the corresponding parameter values were (a) reduced by 50% (magenta-$k_{t,ISGRNA}$, green-$k_{c,V}$) and (b) increased by 50% (black-$\mu_{ISGRNA}$, orange-$k_{r,V}$), and blue-$k_{t,V}$).

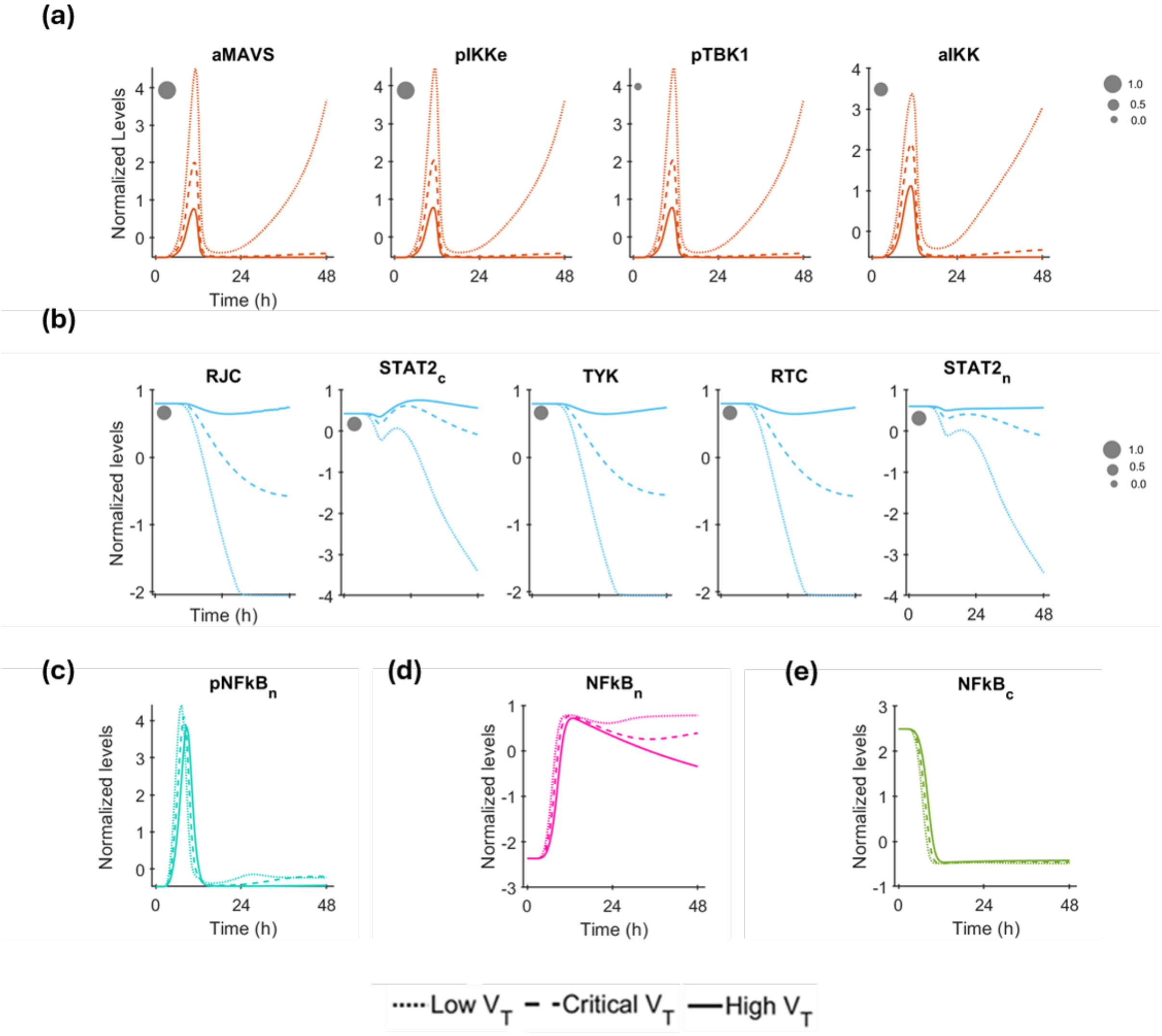


Figure S9: **Normalized dynamics based cluster analysis**
The temporal profiles of components of each cluster (a) Early responders cluster, (b) RTC-TYK cluster, (c) $pNF\kappa B_n$ cluster, (d) $NF\kappa B_n$ cluster, and (e) $NF\kappa B_c$ cluster across the 48-hour time course are plotted. The size of the filled grey circle indicates the correlation between the component and the cluster mean (Figure 3b), with larger markers representing smaller distances. As clusters (c), (d), and (e) contain only one component, correlation is not applicable.

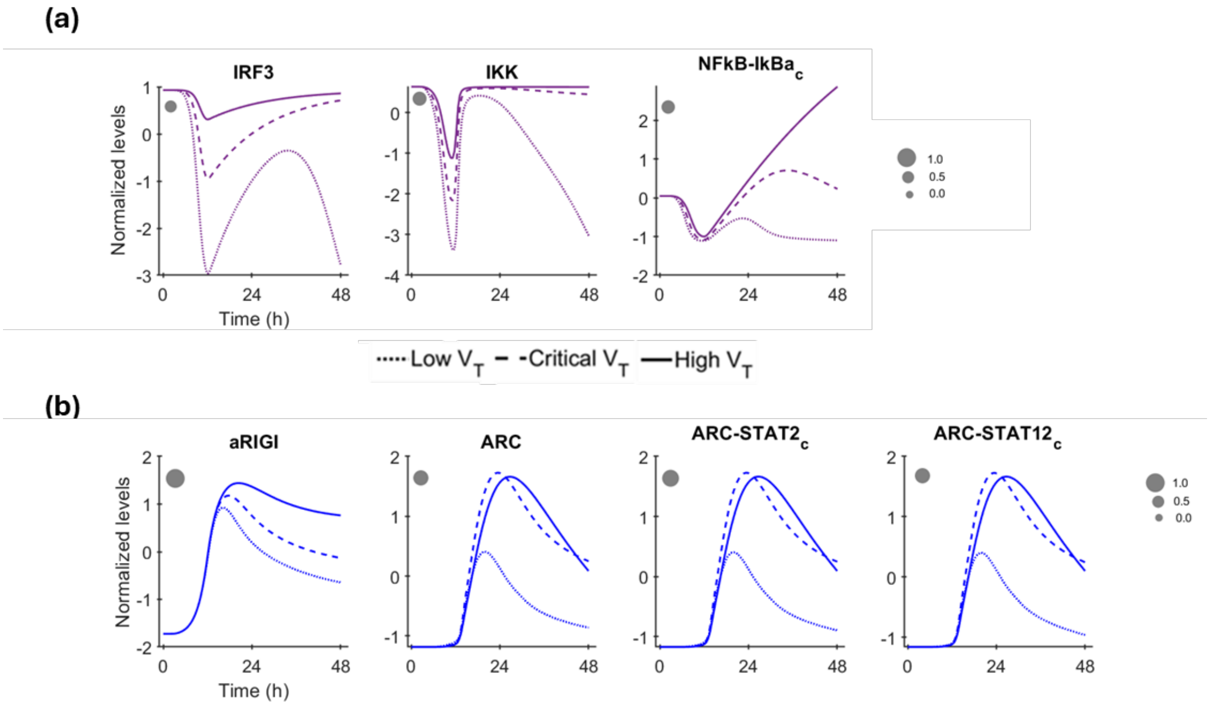


Figure S10: **Normalized dynamics based cluster analysis**
The temporal profiles of components of each cluster (a) IRF3-IKK cluster, (b) secondary responders cluster across the 48-hour time course are plotted. The size of the filled grey circle indicates the correlation between the component and the cluster mean (Figure 3b), with larger markers representing smaller distances.

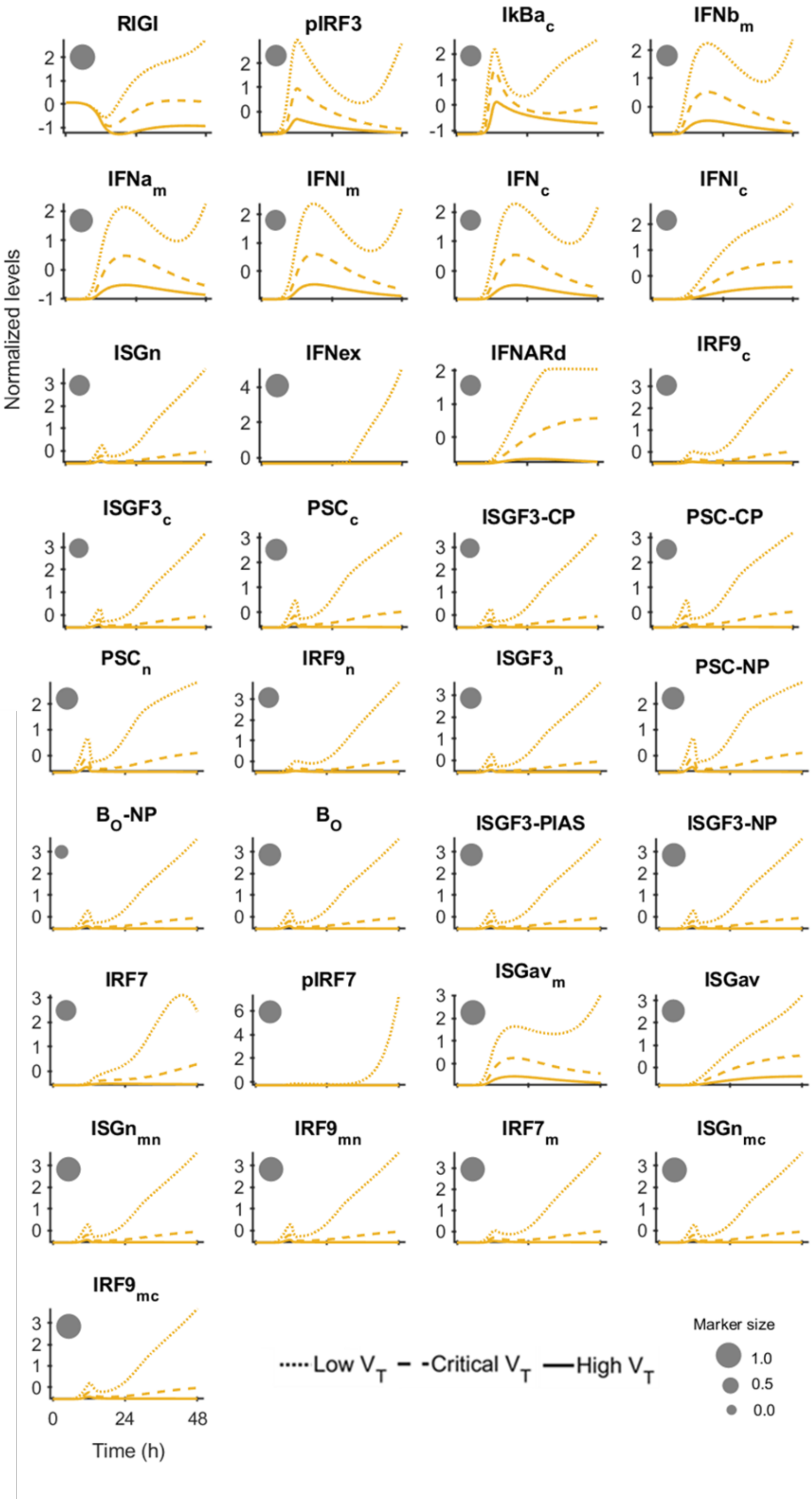


Figure S11: **Normalized dynamics based cluster analysis**

The temporal profiles of components of each cluster IFN-ISG transcriptional hub cluster across the 48-hour time course are plotted. The size of the filled grey circle indicates the correlation between the component and the cluster mean (Figure 3b), with larger markers representing smaller distances.

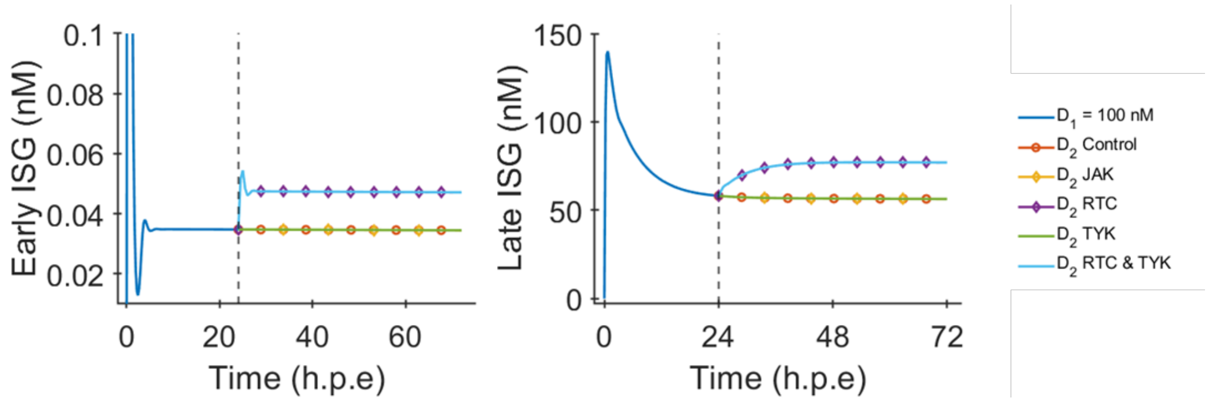


Figure S12: **Dose-dependent desensitization of cellular innate immunity**
Kinetics of (a) an early ISG (ISGn) transcript involved in immune suppression, and (b) a late ISG (ISGav) transcript associated with viral suppression, in response to a two-dose interferon (IFN) treatment. The first IFN dose (100 nM) is administered at 0 h post-exposure (h.p.e.), followed by a second 100 nM dose at 24 h.p.e. The following five conditions are shown: $D_2$ Control (orange circles): Two-dose IFN treatment without any restoration of signaling components. $D_2$ JAK Restored (yellow triangles): JAK levels restored to baseline before the second IFN dose. $D_2$ RTC Restored (purple diamonds): Receptor–TYK2 complex (RTC) levels restored before the second IFN dose. $D_2$ TYK Restored (green line): TYK2 levels restored before the second IFN dose. $D_2$ RTC and TYK Restored (light blue line): Both RTC and TYK2 levels restored before the second IFN dose. Dashed vertical lines indicate the timing of the second IFN dose. These results suggest that restoration of upstream signaling components (particularly RTC) can partially recover ISG responsiveness upon repeated IFN stimulation.

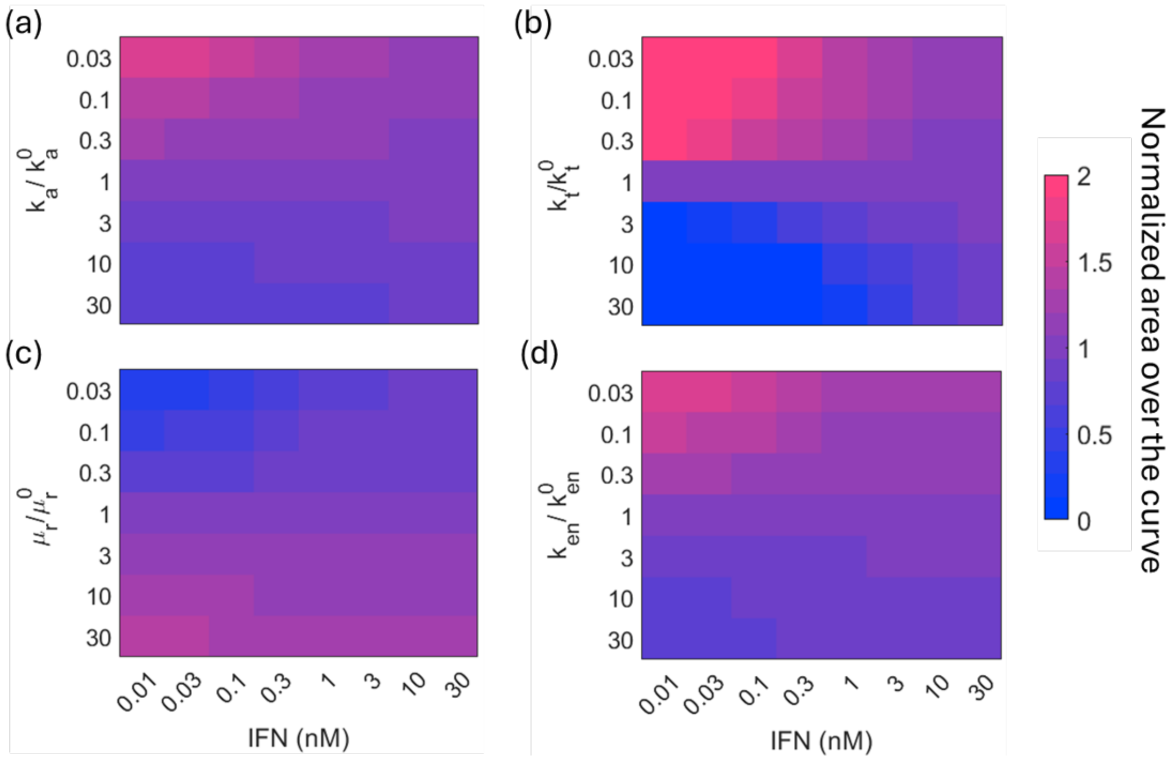


Figure S13: **Exploring IFN: From Preventive Strategies to Therapeutic Interventions**
a) The heatmap shows how the prophylactic efficacy (given by nAOC) varies as a function of IFN concentrations and viral egress rate ($k_{a,V}$). b) The heatmap shows how the prophylactic efficacy (given by nAOC) varies as a function of IFN concentrations and viral entry rate ($k_{en,V}$). c) The heatmap shows how the prophylactic efficacy (given by nAOC) varies as a function of IFN concentrations and viral translation rate ($k_{t,V}$). d) The heatmap shows how the prophylactic efficacy (given by nAOC) varies as a function of IFN concentrations and viral RNA degradation rate ($mu_{r,V}$).
